# A systematic investigation of lactic acid bacteria-derived biosynthetic gene clusters reveals diverse antagonistic bacteriocins in the human microbiome

**DOI:** 10.1101/2022.07.03.498435

**Authors:** Dengwei Zhang, Jian Zhang, Shanthini Kalimuthu, Jing Liu, Zhiman Song, Beibei He, Peiyan Cai, Zheng Zhong, Chenchen Feng, Prasanna Neelakantan, Yong-Xin Li

**Affiliations:** Department of Chemistry and The Swire Institute of Marine Science, The University of Hong Kong, Pokfulam Road, Hong Kong, China; Division of restorative dental sciences, Faculty of Dentistry, The University of Hong Kong, Hong Kong, China; Department of Urology, Huashan Hospital, Fudan University, Shanghai 200040

**Keywords:** Lactic acid bacteria, Biosynthetic gene clusters, Secondary metabolites, Bacteriocins, Human microbiome, Vaginal microbiome

## Abstract

Lactic acid bacteria (LAB) produce various bioactive secondary metabolites (SMs), which endow LAB with a protective role for the host. However, the biosynthetic potentials of LAB-derived SMs remain elusive, particularly in their diversity, abundance, and distribution in the human microbiome. To gain an insight into the biosynthetic capacity of LAB, we analyzed the biosynthetic gene clusters (BGCs) from 31,977 LAB genomes and 748 human microbiome metagenomes, identifying 130,051 BGCs. The found BGCs were clustered into 2,849 gene cluster families (GCFs), most of which are species-specific, niche-specific, and uncharacterized yet. We found that most LAB BGCs encoded bacteriocins with pervasive antagonistic activities predicted by machine learning models, potentially playing protective roles in the human microbiome. Class II bacteriocins, the most abundant LAB SMs, are particularly enriched and predominant in vaginal microbiomes. Together with experimental validation, our metagenomic and metatranscriptomic analysis showed that class II bacteriocins with antagonistic potential might regulate microbial communities in the vagina, thereby contributing to homeostasis. These discoveries of the diverse and prevalent antagonistic SMs are expected to stimulate the mechanism study of LAB’s protective roles in the host and highlight the potential of LAB as a new source of antibacterial SMs.

## Introduction

Lactic acid bacteria (LAB) are a member of Gram-positive, microaerophilic bacteria, which have drawn extensive attention due to their fundamental roles in different biological processes^1^. These bacteria feature lactic acid production in carbohydrate metabolism, which is important in food fermentation. They also contribute to animal production as direct-fed microbes^2^, the chemical industry as cell factories^3^, and the pharmaceutical industry as producers of exopolysaccharides^4^. More importantly, a growing body of evidence reveals the health-promoting effects of consumption of certain LAB strains, rendering them promising candidates for probiotics^5^. Their probiotic actions may arise from multifaceted mechanisms, such as gut microflora regulation, bioactive metabolite production, and immune system modulation, which endow LAB with a protective role for the host^5^.

Despite numerous studies focusing on characterizing LAB as probiotics for microflora regulation, how they impact microbiome homeostasis and host physiology is still not fully understood. Fundamentally, metabolic crosstalk within the kingdoms or with the host is the basis by which microbes, including LAB, engage with microbiome homeostasis. A myriad of metabolites produced by microbes, especially secondary metabolites (e.g. antibiotics and pigments), is among the mediums to achieve a high degree of crosstalk. Secondary metabolite-mediated interactions such as mutualism and antagonism are essential in maintaining microbiome homeostasis^6,7^. Many LAB members, including *Lactobacillus, Streptococcus*, and *Lactococcus*, produce bioactive SMs, ranging from bacteriocins nisin and lactocillin to tetramic acid reutericyclin^8–10^. Bacteriocins are ribosomally synthesized antimicrobial peptides with antibacterial potential and are generally divided into three classes (class I, post-translationally modified peptides (e.g., nisin and lactocillin) ; class II, small unmodified peptides (e.g., amylovorin L and crispacin A); class III, large, heat-labile peptides)^11^. Moreover, bacteriocins, the most extensively studied SMs of LAB, have been recently revealed to play potential roles in shaping the microbiota by modulating microbial composition and inhibiting pathogens^12^. However, these studies of LAB SMs have largely focused on structures, mechanisms of action, or therapeutic potential in preventing infection in a case-by-case manner^13^. The landscape of LAB SMs, particularly their diversity, prevalence, and potential roles in the human microbiome, remains elusive. Therefore, it is still unknown to what extent LAB-derived secondary metabolites (LAB SMs) are actively involved in microbiome homeostasis.

With the development of bioinformatics techniques, the recent explosion of sequenced bacterial genomes and metagenomes provides fresh opportunities for large-scale biosynthetic analysis at both single species and community levels. Here we harnessed recent advances in biosynthetic and metagenomic analysis to investigate the untapped biosynthetic potential of LAB SMs systematically. Leveraging this comprehensive analysis of 31,977 LAB genomes and 748 human microbiome metagenomes, we gained previously undescribed insights into the biosynthetic capacity of LAB SMs and their diversity, abundance, and distribution in the human microbiome. We found that most LAB SMs exhibited pervasive antagonistic activities, exemplified by experimentally validated class II bacteriocins, potentially playing protective roles in the human microbiome. To our best knowledge, this is the first largest survey of LAB biosynthetic potential and the first profile of LAB SMs in the human microbiome. Our findings of the diverse and prevalent antagonistic LAB SMs in the human microbiome, particularly the vaginal microbiome, provide insight into antagonistic interactions linked to microbiome homeostasis and host health.

## Results

### The landscape of SM biosynthetic potential of LAB

Given that environment or foods are possible LAB sources for the gut microbiome, to profile LAB SMs in the human microbiome, we first collected LAB genome data from different sources to comprehensively investigate the biosynthetic potential of LAB SMs. Publicly available bacterial single amplified genomes (SAGs) and metagenome-assembled genomes (MAGs) of LAB were gathered from three databases (RefSeq^14^, PATRIC^15^, and IMG/M^16^) and two previous studies^17,18^, resulting in 40,879 SAGs and 4,575 MAGs in total (Supplementary Table 1). Genomes were then de-duplicated, and their taxonomical classifications were verified and unified using GTDB taxonomy. As a result, 31,977 LAB genomes (27,549 SAGs and 4,428 MAGs, Supplementary Table 2), spanning six families containing 56 genera, were retained for global biosynthetic analysis of LAB SMs (Supplementary Fig. 1a). Using a rule-based BGC detection tool, antiSMASH 6.0^19^, we identified 130,051 BGCs from 30,718 genomes (Fig. 1a, Supplementary Fig. 1b, Supplementary Table 3), including 1,333 nonribosomal peptide synthetase clusters (NRPS, 1.0%), 25,278 polyketides synthase clusters (PKS, 19.7%), 98,810 ribosomally encoded and post-translationally modified peptides (RiPPs, 76.0%), 1,629 terpene (1.3%) and 2,984 BGCs (2.3%) encoding other types of metabolites (Supplementary Fig. 2). The BGCs per genome ranged from 0 to 14, with an average of 4.07. Among the most abundant RiPPs, RiPP-like (formerly annotated as bacteriocin by antiSMASH) topped its list with 72,471 (55.7% of total BGCs). Using BiG-SLiCE^20^ to extract bacteriocin biosynthesis-related domains from 72,471 RiPP-like BGCs (Supplementary Fig. 3 and Supplementary Table 4), we identified 60,497 class II bacteriocins (RiPP-like BGCs that harbor typical class II bacteriocin domains) (46.5%), which are the most abundant LAB SMs. Also worth noting was that 99.8% (25,224/25,278) of PKS were T3PKS (Fig. 1a).

**Fig. 1.**
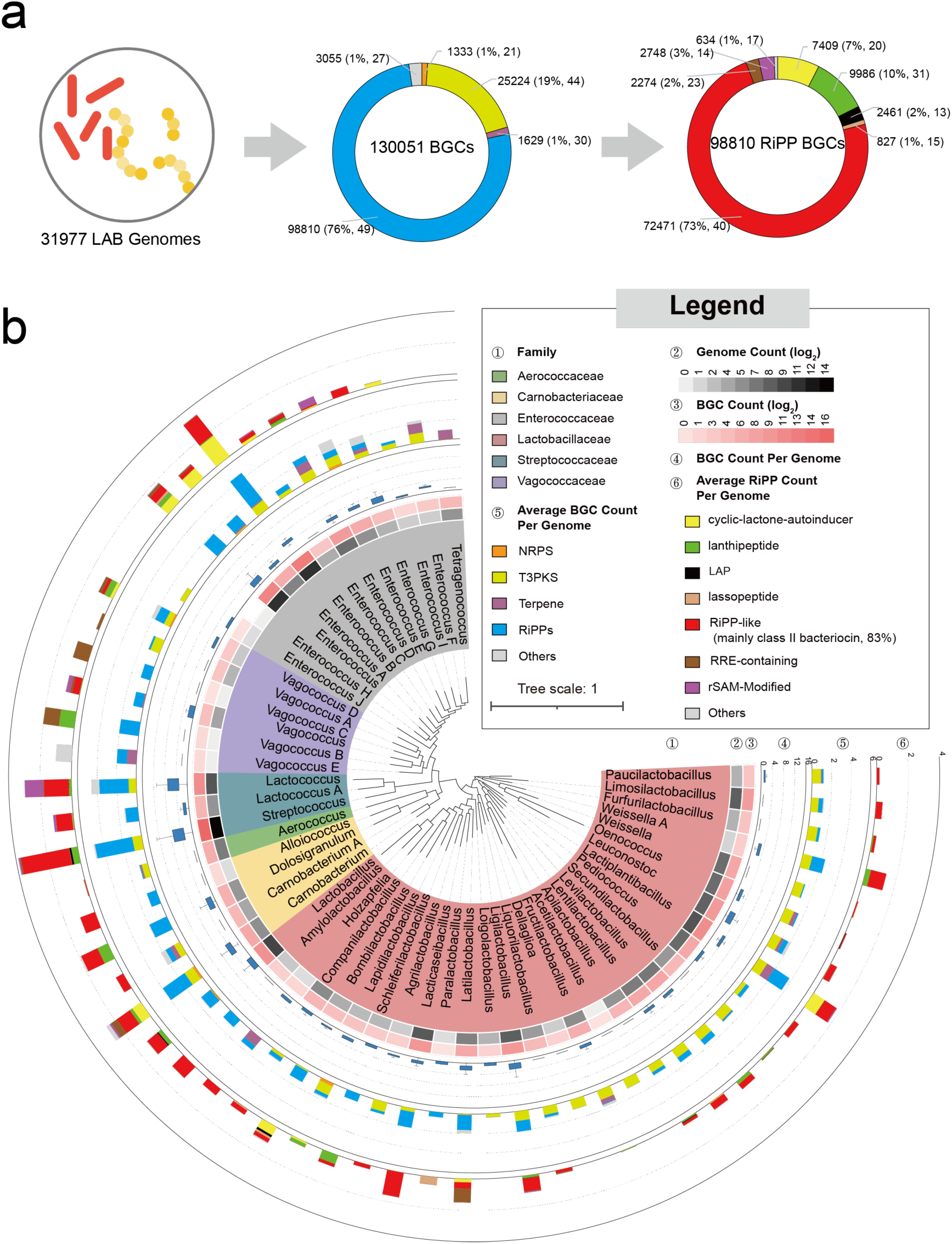
Overview of secondary metabolite biosynthetic capacity in LAB. a, Overall BGCs identified from 31,977 LAB genomes. The numbers outside the brackets indicate BGC count, and numbers represent the corresponding percentage and the count of genera in which BGCs are present. b, Layers are as follows: ①, the maximum likelihood phylogenetic tree based on 120 concatenate marker genes of 56 representative LAB genomes. LAB in this study covers 6 families and 56 genera under GTDB taxonomy; ②, the count of genomes included in this study, with being transformed with log2; ③, log2-transformed BGC count; ④, bar plot showing the BGC count per genome; ⑤, average BGC abundance in LAB genera; ⑥, average RiPPs abundance in LAB genera. In RiPPs term, subtypes not shown here or a combination of >1 subtypes are clustered into “Others”. rSAM-Modified RiPPs consist of RaS-RiPP, ranthipeptide and sactipeptide. No BGCs are detected in one *Enterococcus_H* genome. Figures (a) and (b) share a common legend.

To gain insight into the phylogenetic distribution of BGC in LAB genera, we examined 26,983 BGC-containing SAGs, excluding MAGs due to their incompleteness. The biosynthetic capacity varied considerably at the families, genera, or species level (Fig. 1b, Supplementary Fig. 4). The RiPPs and T3PKS BGCs dominated all LAB genera except for *Tetragenococcus*, while terpene BGCs and NRPS BGCs were sporadically distributed in those genera (Fig. 1, Supplementary Figs. 5, 6). Among 55 genera, we found a median of ≥1 RiPPs per genome in 28 genera and one T3PKS in 33 genera. The BGC distribution pattern in LAB contrasts with a recent study showing the dominance of NRPS in ∼1 M bacterial BGCs^21^. Of note, despite being small in genome size, Streptococcaceae generally harbored more abundant BGCs than other families (Supplementary Fig. 4a), with a median of five BGC per genome, exemplified by genera *Lactococcus* and *Streptococcus*. Contrastingly, 33 genera only harbored a median of <=1 BGC per genome, indicating the limited biosynthetic capacity of the LAB majority (Fig. 1b). By comparing 53 LAB genera and 3,805 non-LAB genera (164,417 genomes, Supplementary Table 5), we found comparatively limited biosynthetic capacity in LAB and a significantly strong correlation (Spearman *rho* = 0.712, *P* < 0.001) between bacterial biosynthetic potential and their genome size (Supplementary Fig. 7). The small genomes and reduced biosynthetic capacities in LAB might correlate with their adaptation to nutritionally-rich niches.

### LAB BGCs are species- or even strain-specific

Although BGCs highly vary in gene content, grouping them into families (GCFs) or clans (GCCs) based on architectural relationships of biosynthetic elements is an effective way to uncover the similarity of their encoding products in terms of the chemical features and biological functions^22^. To gain insight into the novelty and diversity of 130,051 LAB BGCs, we extracted BGC features (biosynthetic domains) using BiG-SLiCE^20^ and grouped them based on an all-to-all cosine distance among BGCs^23^. The 129,878 BGCs with features were classified into 2,849 GCFs and 112 GCCs, with a distance threshold of 0.2 and 0.8, respectively (Fig. 2, Supplementary Fig. 8). We further compared the 112 GCCs to the reference known BGCs described in the ’Minimum Information about a Biosynthetic Gene’ (MIBiG) repository^24^. Notably, only three clans, linear azol(in)e-containing peptides (LAP, GCC_29), RiPP-like (GCC_84), and class II lanthipeptides (GCC_110), were closely similar to known BGCs (average cosine distances < 0.2), leaving the vast majority unknown. This highlights the huge knowledge gap of LAB SMs and demonstrates the potential for discovering novel chemistry from the LAB. Of note, the majority of NRPS (73.2%), terpene (99.3%) and T3PKS (99.3%) were clustered into one respective clan. In contrast, RiPP BGCs, contributing 83 GCCs with 1,818 GCFs (RiPP proportion > 80% in GCCs/GCFs), were highly diverse due to the diversity of their post-translational modification (PTM) enzyme genes and adjacent genes (Fig. 2a). 15 of 23 GCCs that RiPP-like accounted for >80% were class II bacteriocins, contributing 621 GCFs.

**Fig. 2.**
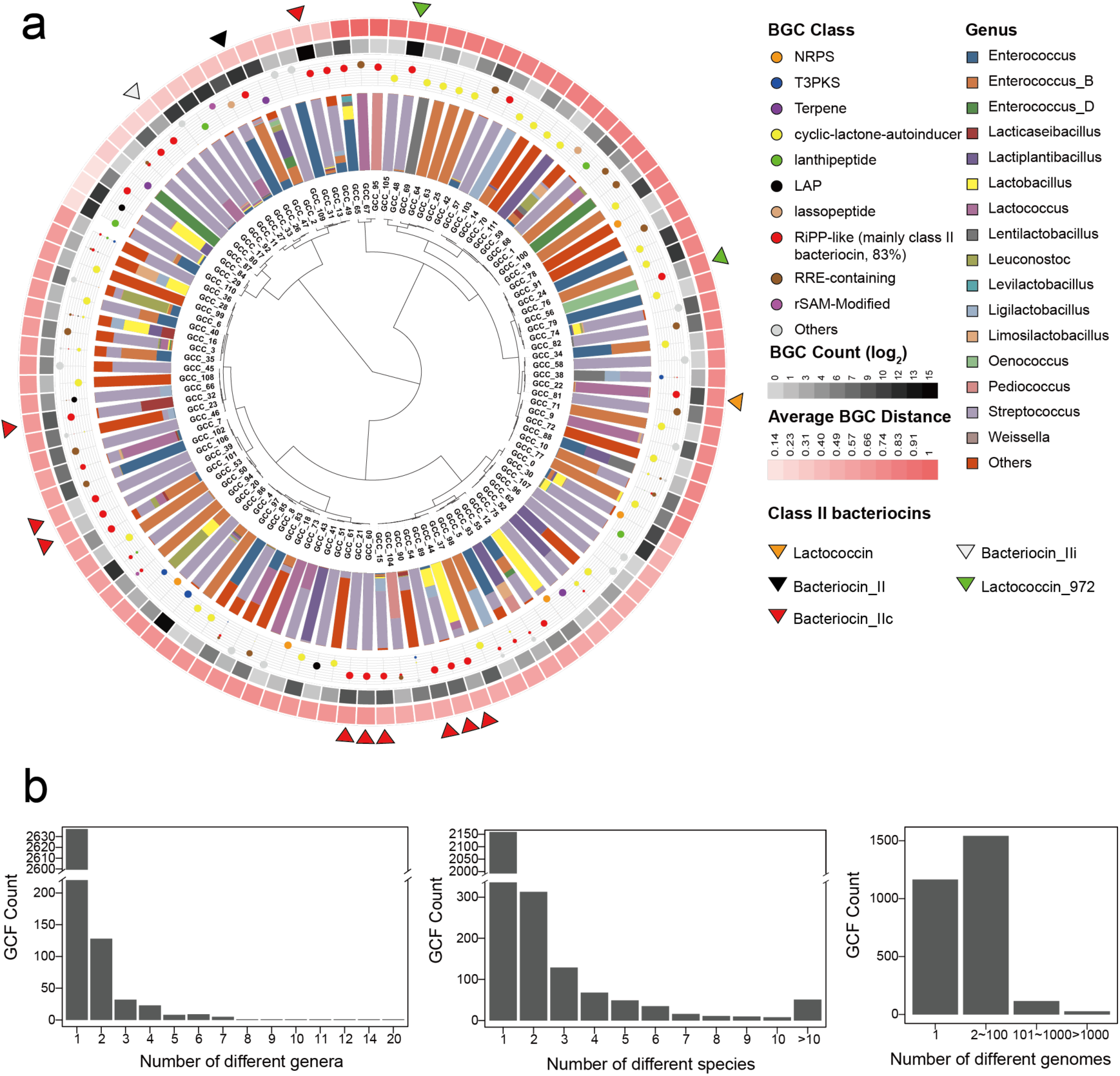
LAB BGCs are diverse and taxa-specific. a, A total of 129,878 BGCs were grouped into 2,849 GCFs and 112 GCCs. The innermost dendrogram is the hierarchical clustering of 112 GCCs, on the basis of their average cosine distance to MIBiG BGCs. The next outer layer is the proportion of different genera. Genera with genome count < 200 are grouped into “Others”. The following layer is the proportion of BGC classes, which is proportionate to point size. The two outer layers refer to log2-transformed BGC count and average distance to MiBiG BGCs. The triangles denote clans dominated by particular class II bacteriocins-related domains (proportion > 80% in one clan). The predominant bacteriocins-related domains are shown in Supplementary Fig. 10. b, The bar plot shows the number of GCFs present in different genera (left), species (medial), and genomes (right).

In prokaryotic genome evolution, the conserved genes cross genera are more likely to contribute to essential ecological processes, whereas species- or even strain-specific genes often arise from natural selection, thus enhancing niche adaptation or host fitness^25^. In this context, we next examined the distribution and diversity of genus- or species-specific BGCs. While GCC clustering shows the distribution and novelty of LAB SMs, a fine resolution of GCF clustering can offer an insight into the diversity of BGCs that are predicted to encode similar natural products. We found that the majority of GCFs were genus-specific (92.6%, 2,637/2,849) and species-specific (75.8%, 2,159/2,849). Remarkably, 1,165 GCFs (40.9%) contained only one BGC harbored by a specific strain (Fig. 2b). In contrast, only 7% (212) were cross-genus GCFs, including 142 RiPPs (present in 2-20 genera), 17 NRPS (in 2-3 genera), 21 T3PKS (in 2-9 genera), and 9 terpenes (in 2-7 genera) (Supplementary Figs. 9, 10). Among these 142 cross-genus RiPP GCFs, 62 are class II bacteriocins. Owing to this high GCF diversity between genera, we did not observe a phylogenetic relationship in GCF presence/absence (Supplementary Fig. 8). Considering the wide presence of LAB in different niches^1^, a high proportion of species- and strain-specific BGCs might be a consequence of niche selection. Of note, the taxa-specificity analysis of GCF may be affected by the imperfect BGC boundary prediction of antiSMASH^26^.

### LAB BGCs are diverse and niche-specific in the human microbiome

Numerous studies have revealed the variable prevalence of LAB species in the human microbiome^18^, raising the question of to what extent their SMs vary in different body sites. Thus, we next explored the profile of LAB BGCs in the healthy human microbiome by re-visiting 748 metagenomes of six body sites from the Human Microbiome Project (HMP)^27^. These sites included aerobic (anterior nares, representing skin microbiome), microaerobic (supragingival plaque, buccal mucosa, tongue dorsum and posterior fornix, representing oral and vaginal microbiome), and anaerobic (stool, representing gut microbiome) environments (Supplementary Table 6). In line with a previous larger-scale study^18^, LAB exhibited variable abundance and prevalence in different body sites (Fig. 3a). Of note, the genus *Lactobacillus* dominated in the vagina with a median abundance of 99.0% [interquartile range (IQR), 91.0%-99.8%], whereas *Streptococcus* was moderately abundant but highly prevalent in six body sites.

**Fig. 3.**
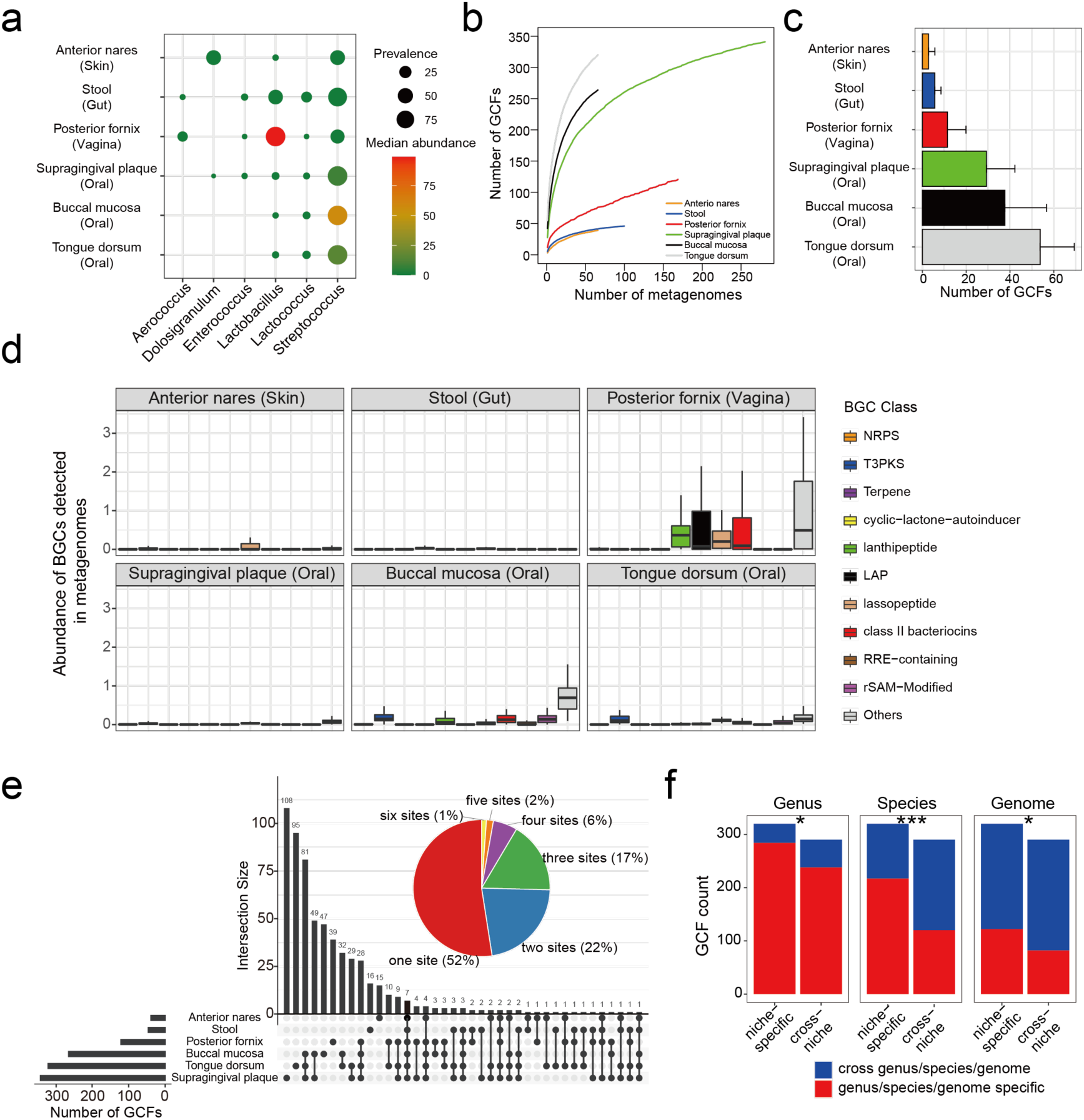
LAB BGCs in the human microbiome are variable and niche-specific. a, The prevalence and abundance of LAB genera in the microbial communities of six body sites. b, GCF accumulation curve, reflecting the number of GCFs detected will increase as more samples are included. c, Number of GCFs detected in six body sites. Data are mean ± standard deviation, with only the upper error bar being shown. d, Boxplot showing the abundance of different BGCs detected in six body sites. Class II bacteriocins here are RiPP-like BGCs that contain class II bacteriocins-related domains identified by BiG-SLiCE. e, The pie chart shows the proportion of 610 GCFs detected in how many sites. Corresponding percentages are shown in the brackets. The bar plot on the left refers to the number of GCFs in each site. The bar plot on the top depicts the number of GCFs of each intersection. Connecting lines are drawn if an intersection is present in more than one site. f, Numbers of genus/species/genome-specific GCFs and cross-genus/species/genome GCFs that were detected in one site (niche-specific) or more than one sites (cross-niche). The Chi-squared test gave *P* values. *, 0.01 < *P* < 0.05; ***, *P* < 0.001.

To profile the diversity, abundance, and distribution of LAB BGCs in the human microbiome, we de-duplicated 130,051 BGCs to 24,222 representative BGCs to map the BGCs upon metagenomes, with a 0.8 cutoff of nucleotide identity to avoid spurious read mapping. The number of LAB BGCs detected in six body sites varied considerably, with the highest in the oral cavity and the lowest in the skin (Supplementary Fig. 11a), probably due to the variable abundance of LAB. The detected 5,687 BGCs belonged to 610 GCFs, including 71 T3PKS and 312 RiPPs with 92 class II bacteriocins (Supplementary Fig. 11b). GCF accumulation curve revealed that more GCFs would be detected in those body sites as more samples were included (Fig. 3b). The three oral sites were the richest in GCFs (averaging 29, 38, and 54). Compared to the oral cavity, the vagina harbors a lower diversity of GCFs (averaging 12) but a significantly higher abundance of LAB BGCs (Fig. 3c, d). Particularly, the vaginal microbiome harbored a high abundance of class II bacteriocins, lassopeptide, lanthipeptide, and LAP. Of note, influenced by sequencing depth, the diversity and abundance of LAB BGCs in the human microbiome are underestimated. Of 610 detected GCFs, ∼52% were niche-specific in one of six sites, which accounted for 18% - 38% of GCFs in a particular site (Fig. 3e). We also observed that those niche-specific GCFs were generally species-specific (Chi-squared test, *P* < 0.001), but not genus-specific (*P* = 0.026) nor strain-specific (*P* = 0.013) (Fig. 3f). This result indicated that niche-specific GCFs were derived from different species residing in distinct niches, which may provide a competitive advantage to the niche adaptation of their hosts. Our genomic and metagenomic analysis of biosynthetic potential revealed that the LAB SMs are diverse and variably prevalent in the human microbiome.

### Most SMs provide LAB with antagonistic potential in the human microbiome

Given the abundance and prevalence of LAB BGCs in the human microbiome, we next want to study the potential bioactivities of BGC-encoding SMs. The bioactivity of SMs encoded by BGCs was recently predicted using machine learning strategies based on chemical fingerprints of predicted compound structure, protein family (PFAM) domains, and other genetic features^28–30^. Here, we adapted four common machine learning classifiers (logistic regression, elastic net regression, random forest, and support vector machines) to predict the bioactivities of LAB-derived SMs. For the training data (950 known BGCs, Supplementary Table 7, Supplementary Fig. 12), ten-fold cross-validation revealed that the random forest classifier outperformed others, with an average area under the receiver operating characteristic curve (AUROC) being 0.76, 0.80 and 0.82, for antibacterial, antifungal, and antitumor or cytotoxic, respectively (Fig. 4a, Supplementary Fig. 13). The performance of the random forest classifier is comparable with previously reported methods^28,29^.

**Fig. 4.**
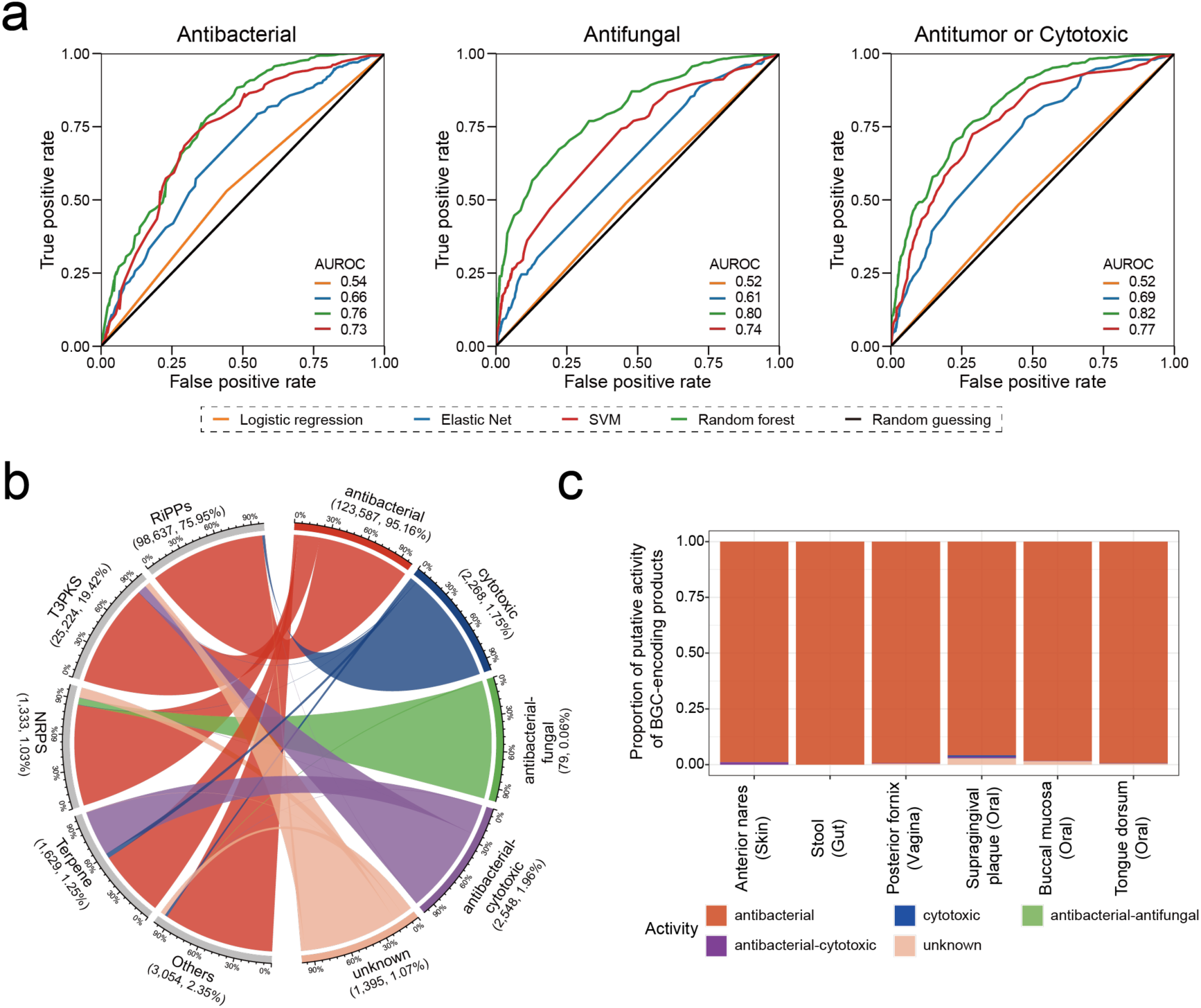
Putative compound activity of LAB BGCs. a, Performance of four machine learning classifiers [logistic regression, elastic net regression, support vector machines (SVM), and random forest] in determining compound activities using 10-fold cross-validation. The receiver operating characteristic (ROC) curves were based on aggregated performances of 10-fold cross-validation. Average AUROC was shown. b, Chord diagram showing the predicted activity of 129,878 BGCs. The scale was the proportion of each BGC class or predicted activity. The number shown in brackets refers to the BGC count and percentage relative to overall BGCs. Antibacterial-antifungal and antibacterial-cytotoxic represent BGCs encoding SMs that were predicted to be both antibacterial and antifungal and both antibacterial and cytotoxic, respectively. c, The proportion of putative activities of BGCs detected in six body sites.

Using the random forest classifier, 129,878 LAB BGCs with features were predicted to encode different bioactive SMs, comprising antibacterial (n=123,587, 95.2%), cytotoxic (n=2,268, 1.8%), antibacterial-antifungal (n=79, 0.1 %), antibacterial-cytotoxic (n=2,548, 2.0%), unknown (1,395, 1.1%). Most BGCs, regardless of BGC classes, were predicted to be antibacterial (Fig. 4b). Of note, 97.8% of RiPPs (96,466/98,637) were predicted to exhibit antibacterial activity, implying that bacteriocins were plenteous in LAB. Most RiPPs with known post-modifications are class I bacteriocins, while most RiPPs-like (83.4%) are class II bacteriocins. Almost 100% of class II bacteriocins (60,494/60,497) were captured as antibacterial SMs, contributing to 48.1% of LAB-derived antibacterial. Those antibacterial BGCs dominated almost all LAB genera (Supplementary Fig. 14), possibly conferring a competitive edge in the microbial community. BGCs encoding putative cytotoxic or antifungal SMs were found in specific species (Supplementary Fig. 15). For example, a certain family of LAP (GCF_199) possibly conferring cytotoxic activity were distributed in ten *Streptococcus* species, especially *Streptococcus pyogenes*, in which common pathogenicity feature endowed by conserved LAP had been reported^31^. In the six body sites, almost all BGCs potentially encoded antimicrobials (Fig. 4c), possibly mediating bacterial antagonism for maintaining microbiome homeostasis. We speculated that LAB harboring antimicrobial SMs, especially class II bacteriocin, could protect the host from pathogen invasion and maintain the microbiome homeostasis via antagonistic interaction^6^.

### Underexplored class II bacteriocins are widely distributed in the human microbiome

The findings of the antagonistic potential of class II bacteriocins and their variable prevalence and predominance in the human microbiome raise the question of what extent the class II bacteriocins may link to the microbiome homeostasis. We next attempted to group class II bacteriocins into subfamily with similar biological functions based on precursor sequence space and investigate their profiling in the human microbiome in detail. To fully reveal the chemical diversity of class II bacteriocin, we first adapted two approaches for the identification of precursor peptides, with hmmsearch^32^ to search Pfam domains of precursors of class II bacteriocin (Supplementary Table 4) and with BAGEL4^33^ which is a tool specifically designed for bacteriocin mining. Combining two approaches, we identified 187,649 precursors from class II bacteriocin BGCs (Fig. 5a, Supplementary Table 8). We then grouped 187,649 putative precursors into 2,005 clusters with a threshold of 50% sequence identity (Supplementary Fig. 16). The sequence lengths of those representative precursors were approximately normally distributed, with a center of ∼55 amino acids (Fig. 5b). The accumulation curve showed that the precursor diversity increased with the number of genomes included, indicating more class II bacteriocins will be disclosed with more genomes sequenced (Fig. 5c). Moreover, we found that only 188 clusters were similar to 333 known class II bacteriocins (identity>90%, coverage>95%), leaving the vast majority (1,817/2,005) underexplored (Supplementary Table 9). Of note, while the rule-based method hmmsearch and BAGEL4 enable a high likelihood of positive detection, at the same time, they probably underestimate the real biosynthetic potentials of bacteriocins.

**Fig. 5.**
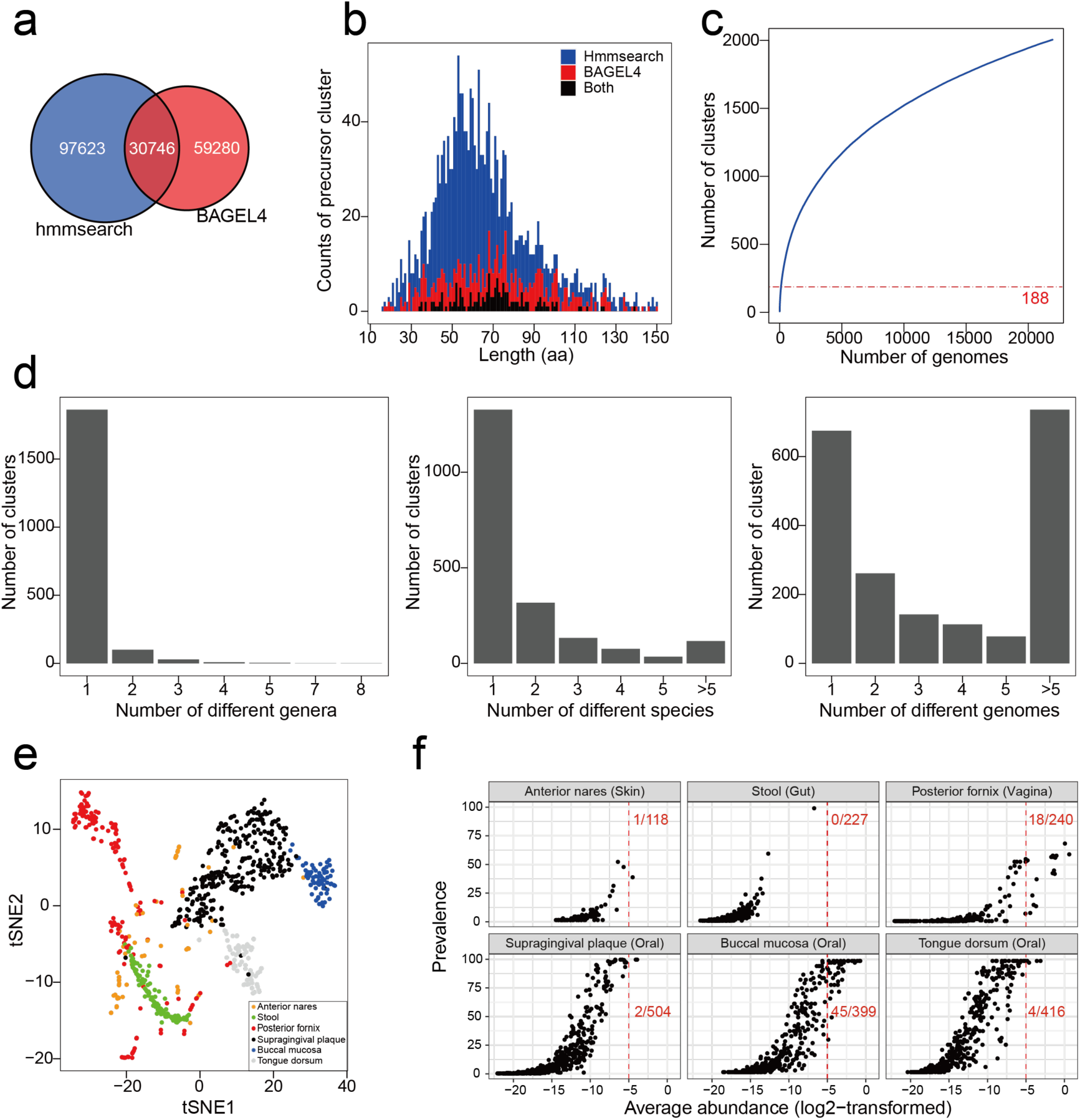
Class II bacteriocins are structurally diverse and variably prevalent in human microbiome. a, The number of putative precursors of class II bacteriocins detected by hmmsearch and BAGEL4. They identified 128,369 and 90,026 putative precursors, respectively, with 30,746 sequences in common. b, Distribution of the length of 2,005 representative precursors, which Cd-hit designated. c, Rarefaction curve of clusters of class II bacteriocin precursors. The Red line shows that 1,775 sequences belonging to 188 clusters were highly similar to known class II bacteriocins (identity>90%, coverage>95%). d, Number of clusters in different genera (left), species (middle), and genomes (right). e, t-SNE plot reveal the distinct profile of class II bacteriocins in different body sites. The posterior fornix (vagina) and anterior nares (skin) show great individual variations, whereas the class II bacteriocins in other body sites are relatively conserved with being clustered together. Each dot represents one metagenome sample. f, The prevalence and average abundance of 644 precursor clusters detected in six body sites. Each dot denotes one precursor cluster. The abundance in the individual is shown in Supplementary Fig. 17. The red numbers are the number of precursor clusters with a log2-transformed average abundance > -5 (shown in red dash line) and the number of precursor clusters detected in each body site.

In line with the taxa-specificity of the GCFs, most class II bacteriocin precursors family were genus-specific (n=1,862, 92.9%) and species-specific (n=1,327, 66.2%), with 33.7% of precursor family being even strain-specific (n=675) (Fig. 5d). We next examined their profiling in the human microbiome. Of 644 clusters detected in six body sites, about 31.8% of clusters were niche-specific in one of six sites (Supplementary Fig. 17a). Moreover, the profiles of class II bacteriocins in different body sites were distinct, with those in the vagina and skin being varied noticeably between individuals (Fig. 5e). Those class II bacteriocins were sporadically present in the skin and gut, whereas some class II bacteriocins were particularly enriched in the oral cavity and vagina with a high prevalence and abundance (Fig. 5f, Supplementary Fig. 17b). Considering the vagina has simple communities with the lowest alpha diversity than other body sites^34^, we reasoned that those enriched and predominant class II bacteriocins might play prominent roles in regulating microbial community in the vagina.

### Keystone class II bacteriocins potentially contribute to vaginal microbiome homeostasis

To examine the class II bacteriocins that may account for the homeostasis of the vaginal microbiome, we constructed the association network between class II bacteriocins and bacterial species at the metagenomic level. We found 23 precursor clusters correlated negatively with various species, indicating their antagonistic potential in regulating the vaginal microbiome (Fig. 6a). Of 23 precursor clusters, 21 clusters were found to be inversely correlated to the Shannon index (Spearman *rho* < -0.4, adjusted *P* < 0.05) (Fig. 6b), further supporting their regulative role in shaping the microbiome. To confirm whether these antagonistic bacteriocins are biologically functional, we next inspected their expression profile in the 180 metatranscriptomic datasets (Supplementary Table 6) and found that most of them were actively transcribed in the vaginal microbiome of healthy individuals (Fig. 6c). Of 21 clusters, three belonged to known class II bacteriocins, including Amylovorin L (cluster_342 and cluster_346, a two-component class IIb bacteriocin) from *Lactobacillus amylovorus* DCE 471^35^ and gassericin T (cluster_94) from *Lactobacillus gasseri* SBT2055^36^, supporting the effectiveness of our association analysis in discovering the regulatory bacteriocins in the microbiome. The other 18 clusters were uncharacterized class II bacteriocins that were also actively transcribed in the vagina.

**Fig. 6.**
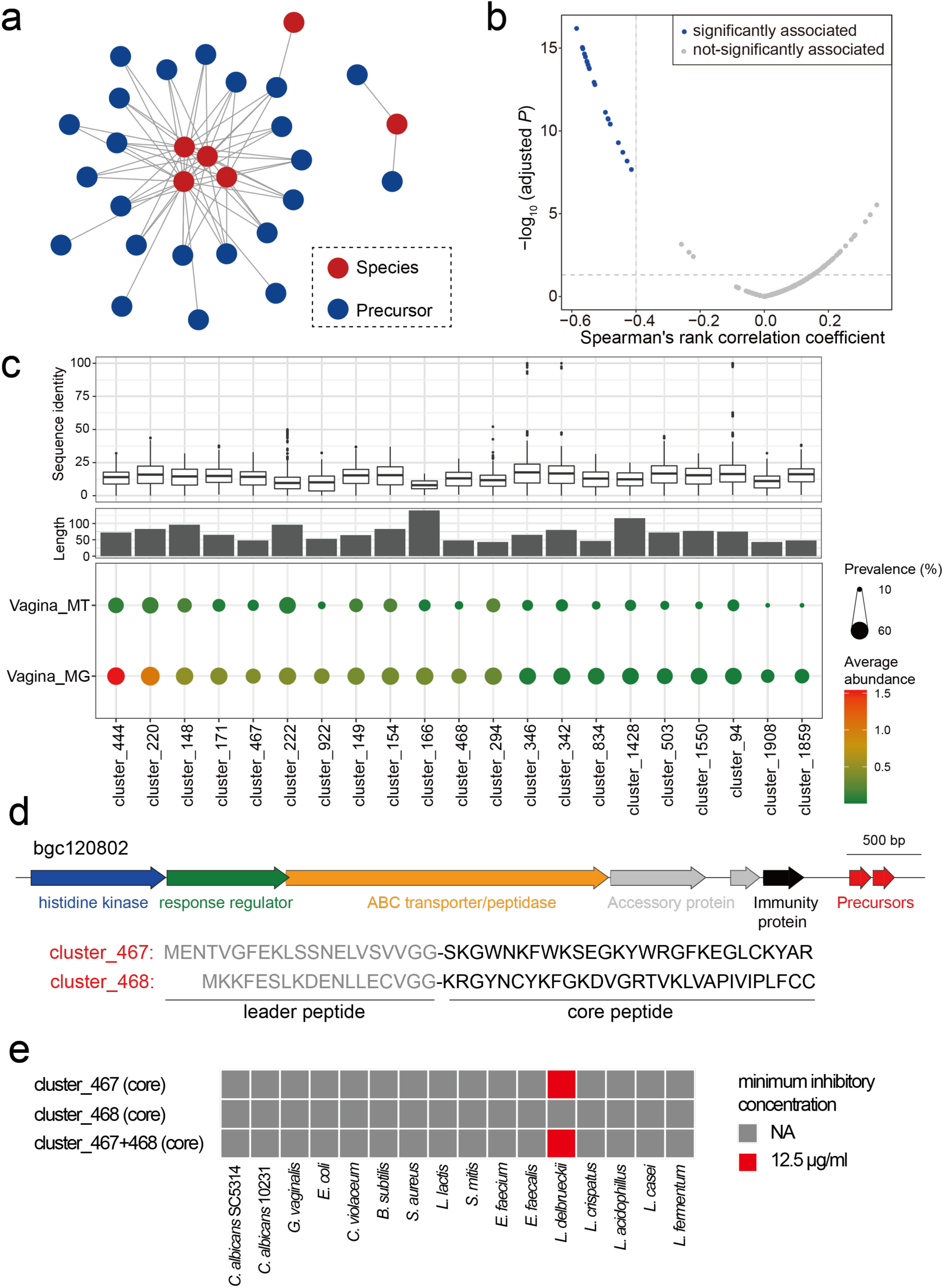
Antagonistic class II bacteriocins potentially play a regulating roles in the vagina microbiome. a, Co-occurrence network of negatively correlated precursor clusters of class II bacteriocins and bacterial species in the vaginal microbiome. 23 clusters are negatively correlated with species in community level, with spearman’s rho < -0.3 and adjusted *P* < 0.05 shown in the network. b, Spearman correlation between precursor clusters and alpha diversity (Shannon index). The dash lines denote the cutoff of correlation coefficient < -0.4 and adjust *P* < 0.05. Points refer to 240 precursor clusters detected in the vaginal metagenomes, 21 of which were significantly associated with the alpha diversity of the vaginal microbiome. c, Global sequence identity to known class II bacteriocins (upper), sequence length (middle), abundance and prevalence of 21 clusters (bottom) in the vaginal metagenome (MG, n=169) and metatranscriptome (MT, n=180). d, Gene organization of bgc120802 and sequences of cluster_467 and cluster_468. Putative double-glycines leader peptides are in grey. The bgc120802 harbors specific class II bacteriocins-related genes, including a two-component regulator system (histidine kinase and response regulator), ABC transporter, and immunity protein. e, The minimum inhibitory concentration of core peptides of two precursors. NA: not available, no inhibitory effect detected with 200 μg/ml.

We next sought to validate the biological functions of those putative bacteriocins experimentally. For proof of principle, we selected two precursor clusters (cluster_467 and cluster_468) with high abundance and a short peptide length that make their chemical synthesis practical. Those two precursors were located on BGCs identified from 16 genomes of *L. crispatus* (Supplementary Fig. 18). The precursor sequences in each cluster were identical and featured a canonical double-glycine leader (Fig. 6d, Supplementary Fig. 18). We thus synthesized the core peptides (Supplementary Fig. 19), namely crispacin 467 (27 amino acids) and crispacin 468 (30 amino acids), respectively, and validated their antagonistic activity toward bacteria and fungi. The antimicrobial assay showed that the crispacin 467 exhibited a narrow-spectrum antibacterial activity against phylogenetical-closely related strain *L. delbrueckii* subsp. *bulgaricus* with a minimum inhibitory concentration of 12.5 μg/mL. While inhibitory effects of crispacin 468 were not observed, nor a synergy of them (Fig. 6e). Of note, the crispacin 468 might exhibit antimicrobial activity against other species beyond the tested strains. Taken together, we believe that the bacteriocin producers arm the vagina microbiome with diverse antagonistic bacteriocins, preventing pathogen invasion and stabilizing the microbial community.

## Discussion

Despite increasing evidence revealing the health-promoting effects of LAB in the human microbiome, how they interplay with other microbes and influence microbiome homeostasis in the human is still understudied. Previous biosynthetic analysis of LAB in a limited dataset focuses on particular species or metabolites, proposing their protective roles to host^37,38^. However, the landscape of LAB SMs, particularly their profiles and potential roles in the human microbiome, remains elusive. In this study, we conducted a comprehensive BGC analysis for LAB, significantly enhancing our understanding of the diversity and distribution of LAB SMs. In total, we found 2,849 GCFs of 129,878 BGCs from 31,977 LAB genomes, most of which encoded diverse uncharacterized bacteriocins. Most LAB SMs were predicted to be antagonistic using machine learning models, indicating the potential protective roles of LAB SMs for the human microbiome. However, due to limited training data, our machine learning model can only predict limited bioactivities (antibacterial, antifungal, and antitumor), underestimating another biological potential of LAB SMs. Although such evidence does not exclude the possibility of other biological functions, we believe that antagonistic LAB SMs potentially provide competitive edges to their producers and regulate the microbiome community.

We further investigated human metagenomes of six body sites to disclose the LAB BGCs profile of the human microbiome, revealing that the diverse LAB SMs are abundant and variably prevalent in the human microbiome. Class II bacteriocins, the most abundant LAB SMs, were particularly enriched and predominant in the vaginal microbiome. The taxa-specificity of GCFs and their niche specificity in the human microbiome also suggested that the LAB SMs may provide a competitive advantage to the niche adaptation of their producing hosts. In this study, we grouped them into families and clans based on architectural relationships, a well-accepted approach to study the BGC similarity and prioritize novel BGCs for natural product discovery^22,39^. However, the GCF grouping will be impacted by the greedy determination of gene cluster boundaries in antiSMASH^19^. This influence can be reduced by GCF grouping only based on biosynthesis-related domains extracted by BiG-SLiCE^20^. More rational approaches to comparing the BGCs may address this issue in the future.

Applying omics analysis, we underscored 21 class II bacteriocins actively expressed in the vaginal microbiome and negatively correlated with its α-diversity, which may play prominent roles in regulating homeostasis. For proof of principle, we identified a novel class II bacteriocin produced by *L. crispatus*, namely crispacin 467, which exhibited characteristic narrow-spectrum antibacterial activity against phylogenetical-closely related strains. Our results suggested that bacteriocin armed *L. crispatus* with a protective role in vaginal health^40^. Although previous analyses have disclosed the bacteriocin biosynthetic genes and antibacterial activity in *L. crispatus*^41,42^, little is known regarding their antagonistic bacteriocin except crispacin A^43^. A myriad of class II bacteriocins remains elusive, not only in *L. crispatus* but in the entire LAB, providing enormous potential for novel antimicrobials. While the precursors of 21 bacteriocins were widely present and transcribed in the vaginal microbiome, whether they are produced in situ in the vagina still needs to be examined by metabolomics. Meanwhile, whether they are essential for maintaining the homeostasis of the vaginal environment can be explored in the future using mouse models.

In summary, our study provides a global insight into the biosynthetic potentials of LAB SMs and provides a starting point for the genomics-guided discovery of antagonistic SMs that potentially regulate microbiome homeostasis. Class II bacteriocins predominant in vaginal microbiomes but negatively associated with their diversity are experimentally validated to play antagonistic roles in microbial communities. To the best of our knowledge, our study is the first to systematically unveil LAB SM biosynthetic potentials and their profile in the healthy human microbiome. However, the analysis presented here cannot be considered exhaustive. The machine learning strategies employed to predict the bioactivity of SMs remain refined by knowledge accumulation of LAB SMs and their biosynthesis and bioactivity. Additionally, how LAB employ SMs to shape their microbiome communities in the human niche remains to be studied. Nevertheless, our discovery of the diverse and prevalent antagonistic bacteriocins in the human microbiome paves the path for future studies on their regulating roles in microbiome homeostasis. In addition to enhancing our understanding of the profile of LAB SMs and their potential regulating roles in the human microbiome, the discovery of antagonistic bacteriocins opens up exciting opportunities for future research on various probiotic applications of LABs.

## Methods

### Data acquisition

As defined early, lactic acid bacteria include 14 genera, comprising *Lactobacillus*, *Lactococcus*, *Leuconostoc*, *Pediococcus*, *Streptococcus*, *Aerococcus*, *Alloiococcus*, *Carnobacterium*, *Dolosigranulum*, *Enterococcus*, *Oenococcus*, *Tetragenococcus*, *Vagococcus*, and *Weissella*^44^. Particularly, genus *Lactobacillus* has been reclassified recently^45^, extending to 25 genera consisting of *Lactobacillus*, *Paralactobacillus, Amylolactobacillus, Acetilactobacillus*, *Agrilactobacillus*, *Apilactobacillus*, *Bombilactobacillus*, *Companilactobacillus*, *Dellaglioa*, *Fructilactobacillus*, *Furfurilactobacillus*, *Holzapfelia*, *Lacticaseibacillus*, *Lactiplantibacillus*, *Lapidilactobacillus*, *Latilactobacillus*, *Lentilactobacillus*, *Levilactobacillus*, *Ligilactobacillus*, *Limosilactobacillus*, *Liquorilactobacillus*, *Loigolactobacilus, Paucilactobacillus*, *Schleiferilactobacillus*, and *Secundilactobacillus*. Filtered with taxonomy, genomes from these 38 genera were then retrieved from NCBI reference sequences (RefSeq) database^14^ (as of Aug. 2021, including SAGs only), PATRIC database (including SAGs)^15^, IMG/M database (including SAGs and MAGs)^16^. Besides them, genomes from two previous studies focusing on human gut microbiome (including SAGs and MAGs)^17^ and food-originated LAB (including MAGs)^18^ were included as well. To avoid the reference genome redundancy, genomes from RefSeq were compared to themselves as well as those from other sources using Mash v2.3^46^. Genomes with a Mash distance of 0 were considered identical. Only the one with a minimal number of contigs was retained. As potential misclassification might be present, GTDB-Tk v1.7.0^47^ was further used to confirm and unify taxonomic annotation against GTDB-Tk reference data version r202^48^. There is a slight difference between NCBI taxonomy and GTDB taxonomy^49^. Under GTDB taxonomy, five genera are sub-divided: *Carnobacterium*, *Enterococcus*, *Lactococcus*, *Vagococcus*, and *Weissella*. Finally, a total of 56 genera belonging to six families (Lactobacillaceae, Aerococcaceae, Streptococcaceae, Vagococcaceae, Enterococcaceae, and Carnobacteriaceae) were considered as members of LAB in this study. Non-LAB genomes were excluded from further analysis.

### Biosynthetic gene cluster analysis

Biosynthetic gene clusters for each genome were annotated by antiSMASH 6.0^19^ with default parameters. The BGCs of 164,417 non-LAB genomes (3,805 genera, excluding 56 LAB genera mentioned above) were also annotated by antiSMASH 6.0. Those non-LAB genomes were the intersection between RefSeq and GTDB repository version r202^50^. Only reference genomes from the RefSeq database in GTDB were included, as they are from isolates instead of metagenome-assembled genomes.

### Clustering BGCs into families and clans

BiG-SLiCE^20^, a tool to cluster sizable BGCs, contains two sets of BGC features, biosynthetic-Pfam and sub-Pfam domains. Those BGC features are sufficient to distinguish distinct BGC classes. As a previous study described^23^, all features of LAB BGCs and 1910 experimentally validated BGCs from the MIBiG 2.0 repository were extracted by BiG-SLiCE v1.1.0, and subsequently used to compute all-to-all cosine distances between BGCs using Python suite SciPy version 1.6.2^51^. The cosine distances were next subject to hierarchical clustering with average linkage, grouping BGCs into families (GCFs, distances < 0.2) and clans (GCCs, distances < 0.8) by Python 3.8 with Scikit-learn version 0.24.2^52^.

### Metagenomics and metatranscriptomics analysis

The raw metagenomic sequencing reads of 748 HMP samples^27^ and the raw metatranscriptomic data of 180 vaginal samples^53^ were acquired from NCBI SRA (Sequence Read Archive)^54^ under project accession number PRJNA48479 and PRJNA797778, respectively. Fastp 0.21.1^55^ with default parameters was adopted for detecting and removing low-quality sequencing reads. High-quality metagenomic sequencing reads were subjected to kneaddata (https://github.com/biobakery/kneaddata) for discarding reads belonging to the human host, through searching against the human reference genome (GRCh38.p13) from GENCODE^56^; high-quality metatranscriptomic reads were also subjected to SortMeRNA v4.3.4^57^ for removing reads derived from ribosomal RNAs. Following that, MetaPhlAn v3.0.13^58^ was used for taxonomic profiling. Prior to assessing the abundance of BGCs in metagenomics and metatranscriptomic data, we used a modified script from BiG-MAP^59^ to de-duplicate 130,051 BGCs. To reduce the computational load, we de-duplicated them within each GCF, at a 0.8 nucleotide identity threshold, leading to 24,222 non-redundant BGCs, the nucleotide sequences of which were used to generate the reference database. Next, the non-host metagenomic and metatranscriptomic reads were mapped to this BGC reference using Bowtie 2 v2.3.5.1^60^, with a parameter of “-k 1”. We then utilized featureCounts v2.0.3^61^ (with parameters of “-T 30 -f -p -B -C -t CDS -g ID -M -O -- fracOverlap 0.2”) to assign sequencing reads to the BGC genes. When calculating the abundance of a BGC, we only considered the core and additional biosynthetic genes, excluding the other genes such as transporters, regulators, transposases, and so forth. For each BGC, a corresponding GTF (General Transfer Format) annotation file was generated by antiSMASH. We retrieved the biosynthetic-related genes (tag “biosynthetic” for the core biosynthetic genes and “biosynthetic-additional” for the additional biosynthetic genes) according to the “gene_kind” tag in GTF files. A BGC was considered present in a metagenomic sample when fulfilling the following criteria: (1) the percentage of biosynthetic-related genes detected is over 50% of total biosynthetic-related genes in a BGC; (2) at least one core biosynthetic gene was found in a BGC. The abundance of a BGC was computed via the equation (1):

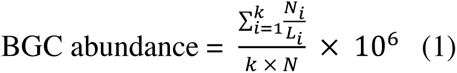

N_i_ represents the number of reads mapped on a biosynthetic-related gene; L_i_ represents the gene length; k represents the number of biosynthetic-related genes in a BGC; N represents the total number of high-quality non-host reads in a metagenome/metatranscriptome sample.

### Prediction of secondary metabolite activity

To predict the activity of BGC-encoding compounds, we used mlr v2.19.0^62^ to perform machine learning. The training dataset comprising 950 MIBiG BGCs with known activities (antibacterial, antifungal, and antitumor or cytotoxic, or other activities) was gathered by Walker *et al.*^29^. BGC features of those known BGCs were extracted by BiG-SLiCE^20^. Prior to training models, we removed BGC features present in < 10 BGCs. Rather than multiclass classification, binary classification was adopted for each activity class since a molecule might have multiple functions. Four two-class classifiers (namely logistic regression, elastic net regression, random forest, and support vector machines) were adopted for binary classification to the activities of BGC products. In order to obtain the honest performance of four classifiers, we measured their accuracies using 10-fold cross-validation. Moreover, 3-fold cross-validation was adopted in each test to tune the hyperparameters, generating 30 instances for each classifier. The average AUROC was used to evaluate the performance of four classifiers. The function *generateThreshVsPerfData* was used to generate data on threshold vs. performances, which was further adapted for plotting the ROC curve. Chord diagram showing the association between BGC classes and predicted activities of their products was plotted using R package *circlize* v0.4.13^63^. Sankey diagram showing the association between species and BGC classes was done by package *networkD3* v0.4^64^.

### Precursor of class II bacteriocins

In order to pinpoint the precursors of class II bacteriocins, we first used Prodigal-short^65^ to identify all small ORFs. We then used hmmsearch^32^ to search class II bacteriocins-related domains (provided in Supplementary Table 4) against ORFs of all RiPP-like BGCs. The hits with a threshold of E-value < 0.01 were considered as the precursors of class II bacteriocins. Meanwhile, BAGEL4^33^ was also adopted for searching class II bacteriocins from RiPP-like BGCs. They detected 128,599 and 90,101 putative precursors respectively, with 30,764 in common. We discard 287 sequences that were larger than 150 AAs, retaining 187,649 sequences for further analysis. Those sequences were then grouped into clusters using Cd-hit^66^, with the parameters of “-n 2 -p 1 -c 0.5 -d 200 -M 50000 -l 5 -s 0.95 –aL 0.95 –g 1”. The sequences with an identity of > 50% will be grouped into one cluster, as proteins with > 50% identity generally share a common function^67^. To collect the known class II bacteriocins, we queried NCBI PubMed with the keyword “class II bacteriocin”. Meanwhile, we also included the sequences gathered by Yi *et al.*^13^ as well as the sequence deposited in the BAGEL4 database^33^. In total, 333 sequences of class II bacteriocins were obtained (Supplementary Table 9). As the curated 333 sequences might be the mature peptides, a local sequence aligner, DIAMOND v2.0.15^68^, was utilized to compare 333 known class II bacteriocins to 187,649 precursor sequences with the parameter of “--id 90 -- query-cover 95 --masking 0”. The known class II bacteriocins showed an alignment of identity > 90% and coverage > 95% with 1,775 precursor sequences belonging to 188 clusters that were thus regarded as homologous. For 21 selected precursor clusters, we identified the global identity relative to the known class II bacteriocins using the Needleman-Wunsch algorithm in the function “needleall” of EMBOSS software package^69^. The alignment of precursors was done by MAFFT v7.490^70^ with the parameter of “--maxiterate 1000 --localpair”, and then was visualized using Jalview software^71^. To conveniently inspect the gene organizations of BGCs harboring precursors of cluster_467 and cluster_468, we adopted BiG-SCAPE^22^ for exploring their architectures.

Precursor abundance in metagenome/metatranscriptome samples was computed via the equation (2):

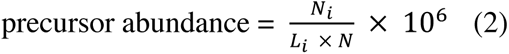

Here, N_i_ represents the number of reads mapped on a precursor gene; L_i_ represents the gene length; N represents the total number of high-quality non-host reads in a metagenome/metatranscriptome sample. The abundance of a precursor cluster is the sum of the abundance of precursors in this cluster.

### Phylogenetic tree construction

GTDB repository version r202^50^ contains 822 representative genomes of 56 LAB genera. The genome with the largest N50 length in each genus was selected as a proxy for its corresponding genus. Consistent with the previous approach to constructing bacterial reference trees^50^, the multiple sequence alignment of the concatenation of 120 phylogenetically informative marker genes of 56 representative genomes was used to infer the phylogenetic tree. IQ-TREE version 2.1.4-beta^72^ was adapted for constructing maximum likelihood (ML) phylogenetic trees, with 1,000 ultrafast bootstrap replicates. In-built ModelFinder^73^ identified the best-fit model as LG+F+R8. Inferred phylogeny was visualized using iTOL^74^.

### Peptide synthesis

The two deducted core peptides of cluster_467 and cluster_468 were chemically synthesized by Sangon Biotech (Shanghai, China). Their molecular weights were confirmed by mass spectrometry, and their required purity was ≥ 90%, which was determined by high-performance liquid chromatography. The synthesized peptide powder was stored at −80 °C and dissolved in sterilized double-distilled water to 2 mg/mL upon use.

### Bacterial and fungal strains

A total of 14 bacterial strains and two fungal strains were used in this study. Their growth conditions are as follows: two bacterial strains (*Escherichia coli* DH5α, *Staphylococcus aureus* B04) were incubated in Luriae–Bertani (LB) culture medium at 37 ℃ under 180 rpm rotation; two strains (*Chromobacterium violaceum*, *Bacillus subtilis* 168) were incubated in LB medium at 30 ℃ with shaking at 180 rpm; *Lactococcus lactis* subsp. *cremoris* MG1363 was grown statically at 30 ℃ in M17 medium; eight bacterial strains (including *Streptococcus mitis*, *Enterococcus faecium*, *Enterococcus faecalis*, *Lactobacillus delbrueckii* subsp. *Bulgaricus*, *Lactobacillus crispatus* ATCC 33820, *Lactobacillus acidophillus*, *Lactobacillus casei*, *Lactobacillus fermentum*) were incubated statically in MRS medium at 37 ℃; *Gardnerella vaginalis* was maintained in the Colombia blood agar at 37C in anaerobic conditions, and the suspension culture is grown by taking a loop full of colonies from the agar plate and incubated in Brain heart infusion broth (BHI), at 37C in anaerobic conditions; two fungal strains (*Candida albicans* SC5314, *Candida albicans* ATCC 10231) were grown in RPMI media at 37 ℃ with shaking at 150 rpm.

### Determination of minimum inhibitory concentrations

The minimum inhibitory concentrations (MICs) of two peptides [individually and in combination (1:1 ration)] against bacterial and fungal strains were performed by broth microdilution. Tested bacterial strains were inoculated overnight in the corresponding culture medium (LB, M17, BHI, or MRS) and at respective growth conditions. The optical density at 600 nm (OD_600_) of bacterial cultures was determined to estimate the bacterial concentration. The bacteria cultures were diluted to ∼5 × 10^5^ CFU/mL using the respective broth. 100 μL aliquots of bacterial suspensions were transferred into 96-well plates containing two-fold serial dilutions of peptides (ranging from 200 μg/ml to 0.19 μg/ml). After incubating for 24 h, bacterial growth was assessed by determining OD_600_. Besides, MIC against the fungal strains was determined according to the CLSI M27-A3 guidelines^75^. Briefly, *C. albicans* strains were cultured overnight in RPMI medium and grown fungal suspensions were centrifuged at 5,000 rpm for 10 min and the pellet was resuspended and washed twice with 1× PBS to remove the dead cells. The fungal inoculum was standardized to 1 × 10^6^ CFU/mL using a spectrophotometer and added to the well plate containing varying concentrations of the peptide (200 μg/mL to 0.19 μg/mL). The media without the peptide serve as a control. The plates were then incubated at 37°C for 24 h, 80 rpm, and the absorbance was measured at 520 nm using SpectraMax 340 tunable microplate reader (Molecular Devices, San Jose, CA, USA). The MIC value was determined as the lowest concentration of the peptides where no bacterial or fungal growth was detected. All assays were conducted in triplicate.

### Statistical analysis and visualization

The accumulations of GCFs detected in metagenomes as well as clusters of class II bacteriocin precursors were computed with function *specaccum* in R package *vegan* v2.5-7^76^. R package *UpSetR* v1.4.0^77^ was adopted for visualizing the intersection of GCFs or precursor clusters detected in different body sites. The alpha diversity (Shannon index) of the vaginal microbiome was calculated with R package *vegan* v2.5-7^76^. To visualize the distribution of class II bacteriocins detected in six body sites, a dimensionality reduction was performed using t-distributed stochastic neighbor embedding (t-SNE), which was done by R package *Rtsne* v0.15^78^. Spearman’s correlations between precursor clusters vs. bacterial species and between clusters vs. Shannon index were computed with the function *corr.test* in R package *psych* v2.1.9^79^, and *P* values were adjusted with the “BH” method^80^. The heat maps in this study were plotted using package *pheatmap* v1.0.12^81^. Cytoscape 3.9.0^82^ was used to visualize the network of similarity of class II bacteriocins and the network of species-precursor correlation. Without a specific statement, other figures were generated using ggplot2 v3.3.5^83^. All statistical analyses were finished in R v4.1.2.

## Data availability

The bacterial genomes are publicly available in NCBI Assembly RefSeq database (https://www.ncbi.nlm.nih.gov/assembly), PATRIC database (https://docs.patricbrc.org/user_guides/ftp.html), and IMG/M database (https://img.jgi.doe.gov/cgi-bin/m/main.cgi). The genomes from human gut are available in the European Nucleotide Archive under study accession ERP116715, and genomes from food metagenomes are available at http://www.tfm. unina.it/DATA001-2020-Pasolli. Genomes can be obtained through the accession numbers provided in Supplementary Tables 1 and 5. The raw data for HMP metagenomes and vaginal metatranscriptomes are deposited in NCBI-SRA under the BioProjects PRJNA48479 (https://www.ncbi.nlm.nih.gov/bioproject/48479) and PRJNA797778 (https://www.ncbi.nlm.nih.gov/bioproject/?term=PRJNA797778), respectively. The samples used in this study are provided in Supplementary Table 6.

## Acknowledgments

This work is partially funded by a Shenzhen Basic Research General Programme (JCYJ20210324122211031) and two Hong Kong Research Grants Council General research grants (HKU27107320 and HKU17115322). The authors would like to thank Dr. Mingqiang Qiao and Wanjin Qiao at Nankai University for providing *Lactobacillus lactis* strain.

## Author contributions

Y.-X.L. and D.Z. conceived of the study, participated in its design and coordination, and drafted the manuscript. D.Z. and J.Z. gathered publicly available data used in this study. D.Z. and S.K. performed MIC determination. J.L., Z.S., B.H., and P.C. performed bacterial culture, compound isolation, and metabolic analysis. D.Z., J.Z., Z.Z., and Y.-X.L. performed data analysis and interpretation. C.F. and P.N. provided advice. Y.-X.L. was involved in the overall supervision of the project. All authors read, revised, and approved the final manuscript.

## Supplementary Figures

**Supplementary Figure 1.**
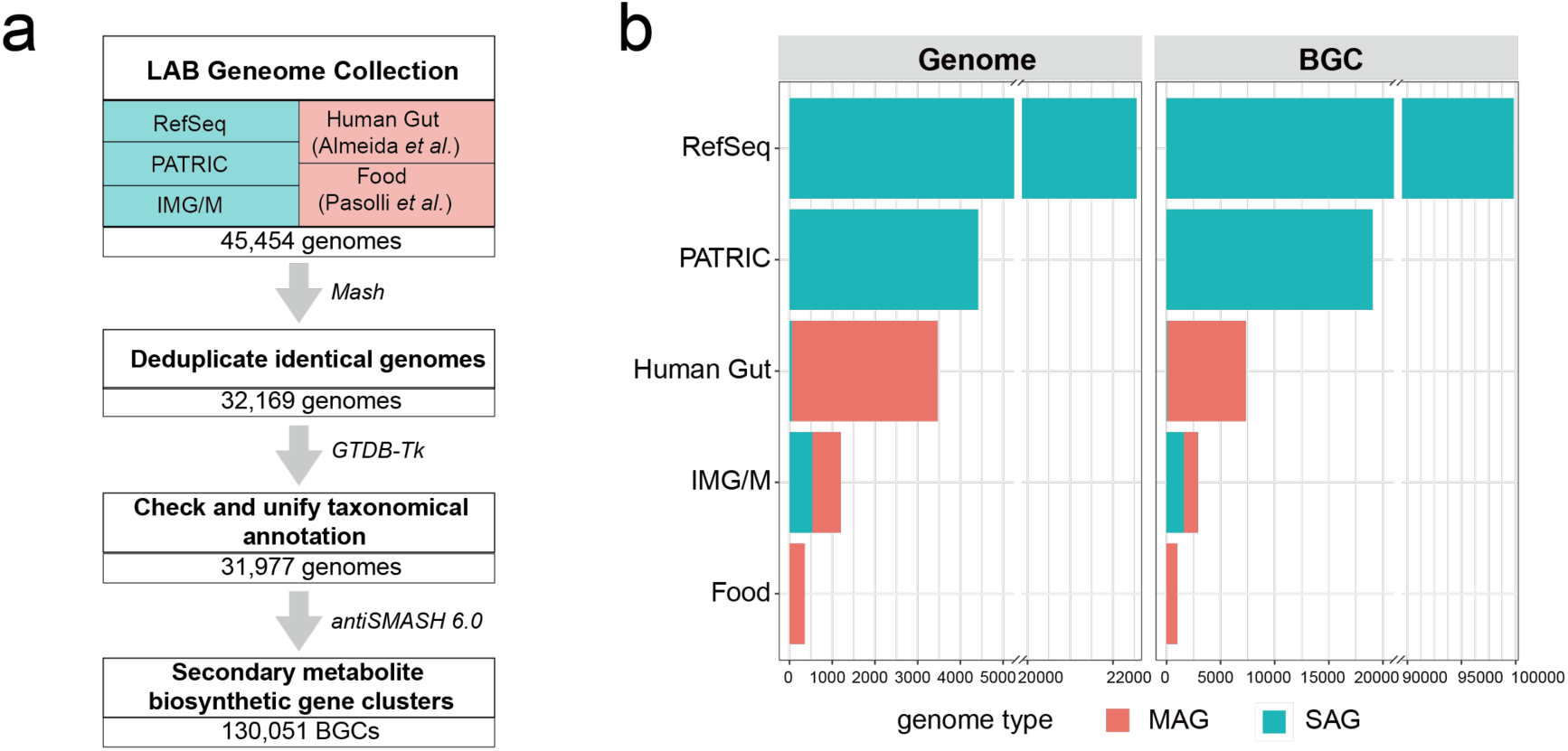
Overview of the data processing. **a**, A flowchart showing the data collection and BGC annotation. LAB genomes were gathered from RefSeq, PATRIC, and IMG/M databases as well as two previous studies focusing on human gut and food. **b**, Bar plot showing the number of LAB genomes and BGCs annotated. From 30,718 LAB genomes, a total of 130,051 BGCs were identified, with 120,501 from SAGs and 9,550 from MAGs.

**Supplementary Figure 2.**
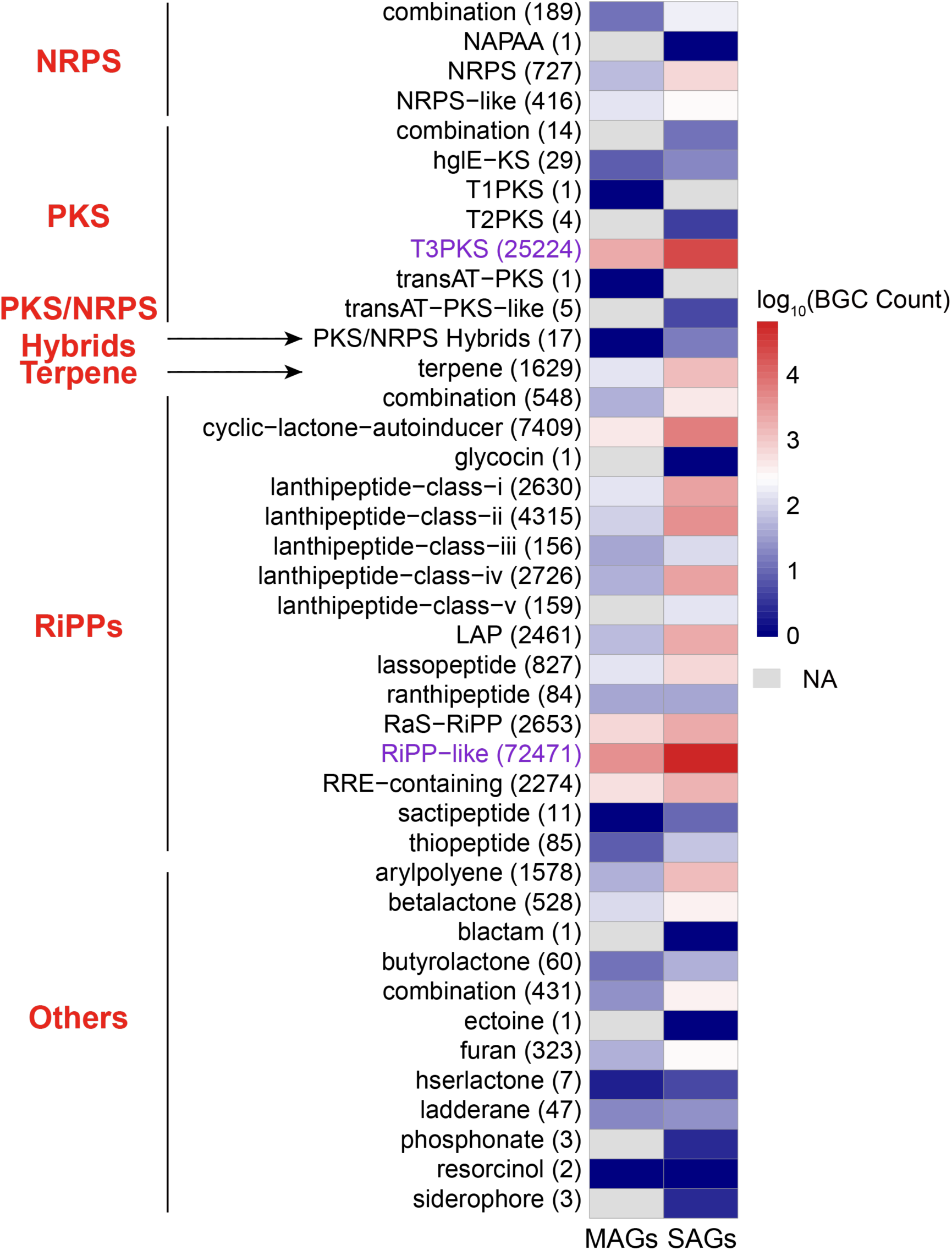
Number of BGCs identified from MAGs and SAGs. The secondary metabolites of BGCs are annotated using antiSMASH. Those BGCs can be grouped into different classes (in red) which are applied in BiG-SCAPE (https://git.wageningenur.nl/medema-group/BiG-SCAPE/-/wikis/BiG-SCAPE%20classes). The numbers in brackets are the sum counts of BGCs from MAGs and SAGs, for each BGC class. The two most abundant BGCs, T3PKS and RiPP-like, are highlighted in purple.

**Supplementary Figure 3.**
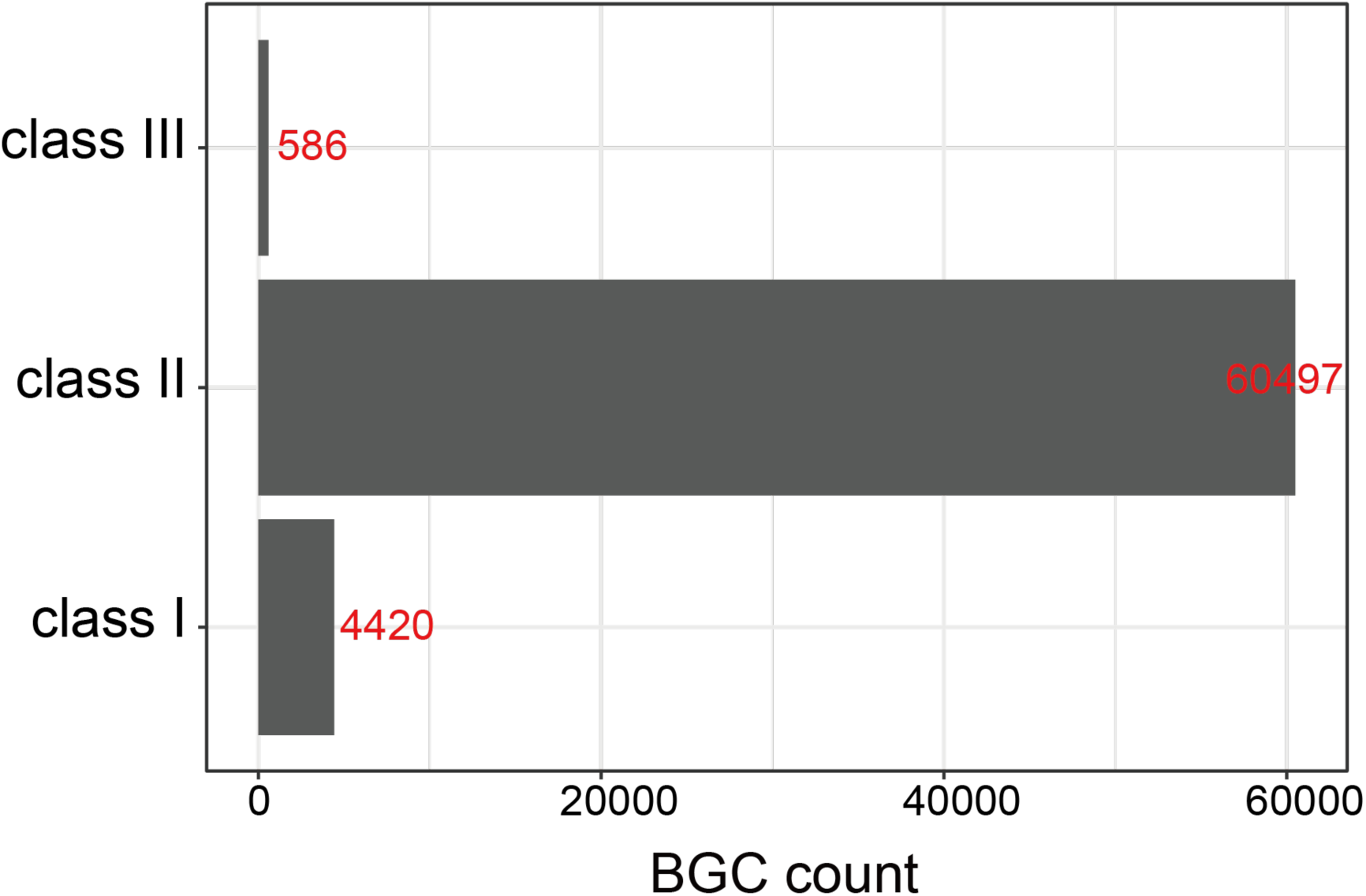
Number of bacterioncins identified from 72,471 RiPP-like BGCs. As antiSMASH will annotate partial class I, class II, and class III bacteriocins as RiPP-like, we then used BiG-SLiCE to extract bacteriocin biosynthesis-related domains (provided in Supplementary Table 4). By this, we further identified 4,420 class I bacteriocins, 60,497 class II bacteriocins (46.5%), and 586 class III bacteriocins (0.5%) from RiPP-like BGCs. It should be noted that other RiPPs such as lanthipeptides, thiopeptides, and lassopeptides could be classed into class I bacteriocins if they were active against bacteria.

**Supplementary Figure 4.**
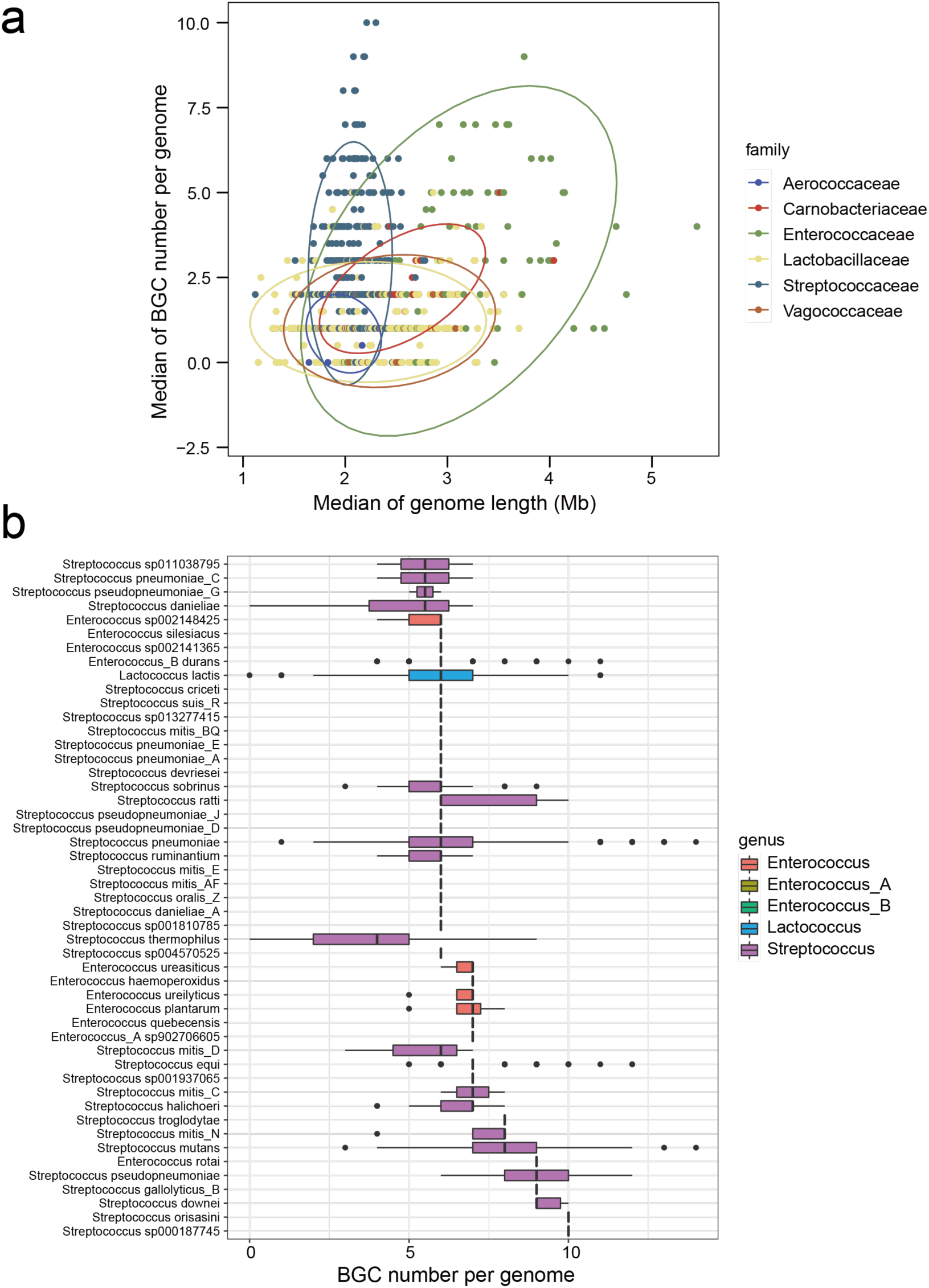
Biosynthetic potential in LAB species. **a**, Scatterplot showing the profile of BGC number per genome within LAB species, colored according to the family level. Each dot stands for one species. Despite being small in genome size, family Streptococcaceae generally harbored more abundant BGCs compared to other families. **b**, The distribution of BGC number per genome within 49 species that have a median number of >5. Of 49 species, the species number belonging to genus *Streptococcus* and *Enterococcus* were 37 and 9, respectively.

**Supplementary Figure 5.**
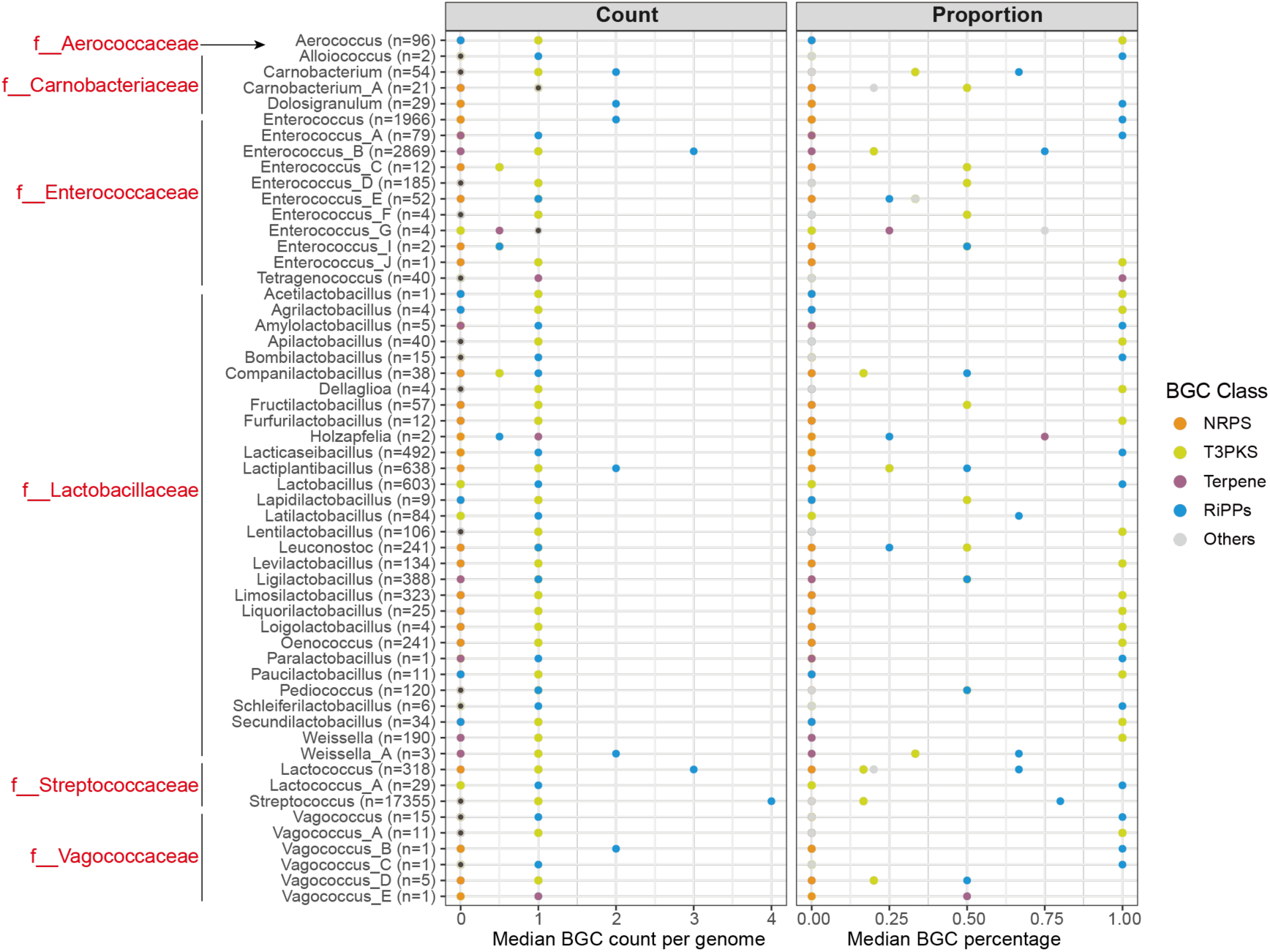
Median BGC count and proportion in SAGs. The numbers in brackets are counts of BGC-containing SAGs for each genus.

**Supplementary Figure 6.**
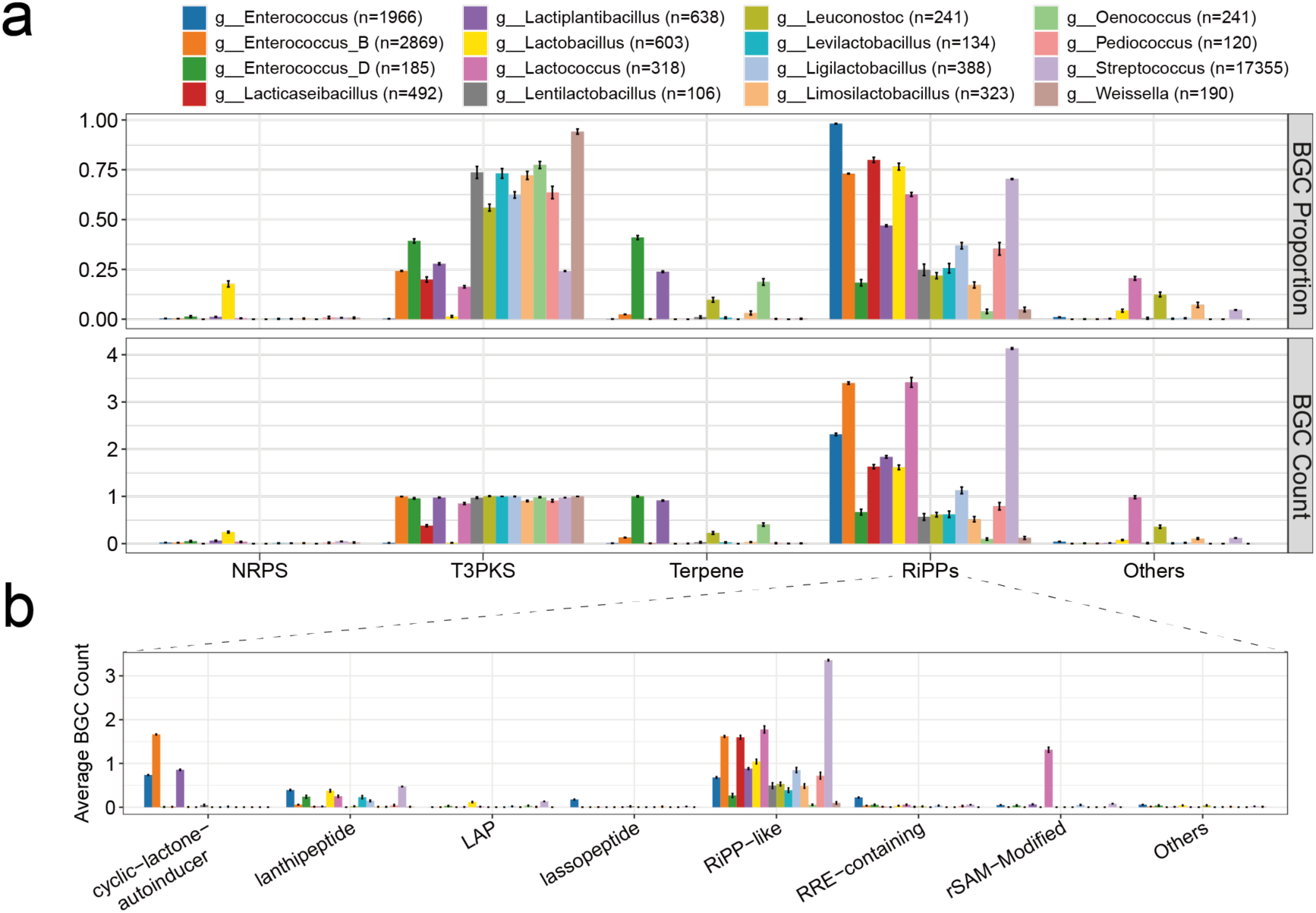
Comparison of BGC proportion and counts in LAB. **a**, Bar plot showing the average BGC percentage (upper) and average BGC count per genome (lower) in 16 genera that comprise over 100 SAGs. To reduce the sampling bise, the average BGC percentage/count was calculated from half of the genomes of this genus, with 1,000 replicates of sampling. The numbers in brackets indicate the number of SAGs for each genus. **b**, The abundance of distinct RiPPs in 16 genera. rSAM-Modified RiPPs consist of RaS-RiPP, ranthipeptide and sactipeptide. Other types of RiPPs or combinations of different RiPPs types are grouped into “Others”. Data are mean ± standard deviation.

**Supplementary Figure 7.**
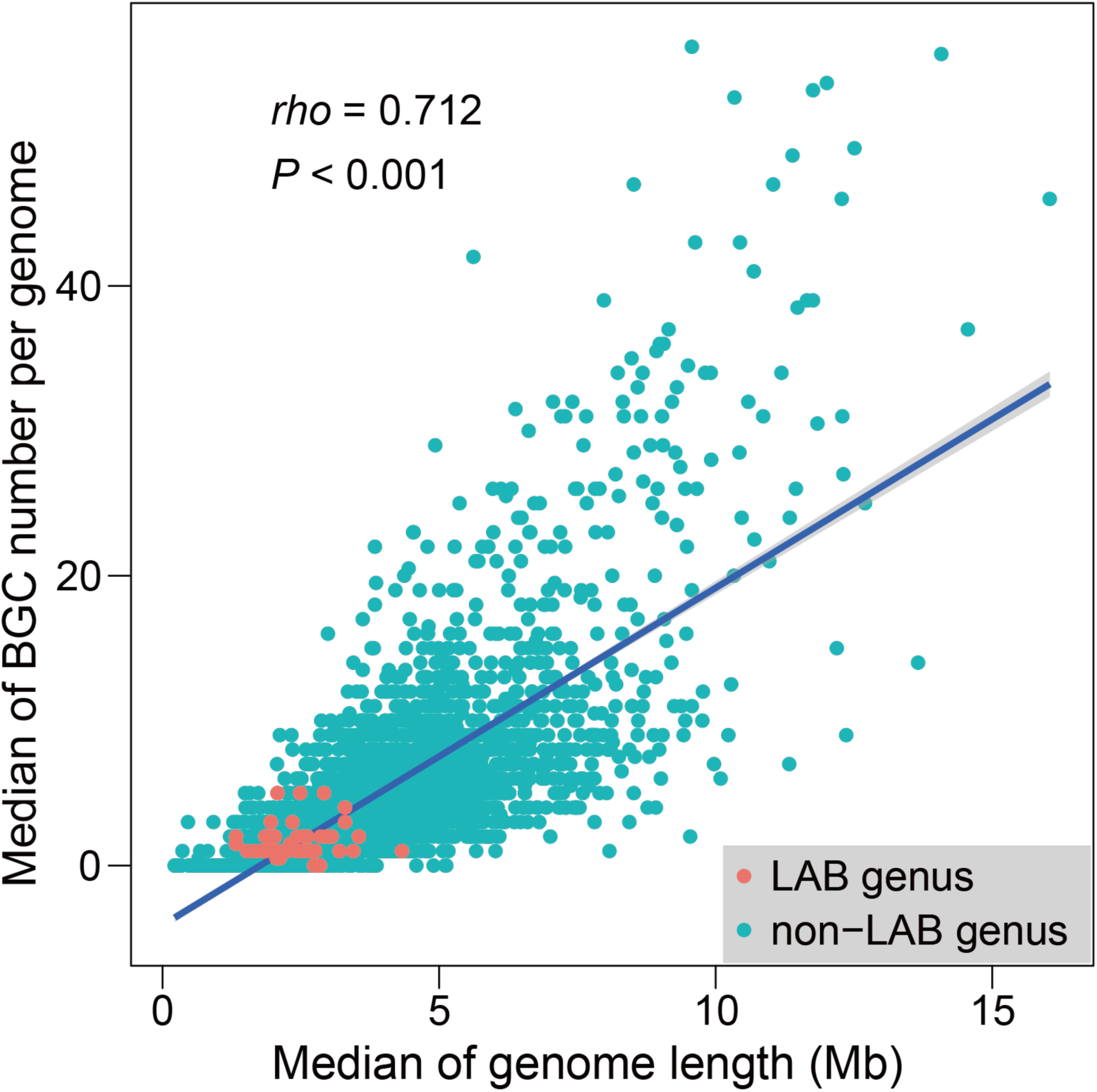
Comparison of SM BGC capacity between LAB and non-LAB genera. Each dot represents one genus, with 56 LAB genera and 3,805 non-LAB genera. Spearman’s rank correlation was carried out, and significance was verified. The grey shade shows the 95% confidence interval.

**Supplementary Figure 8.**
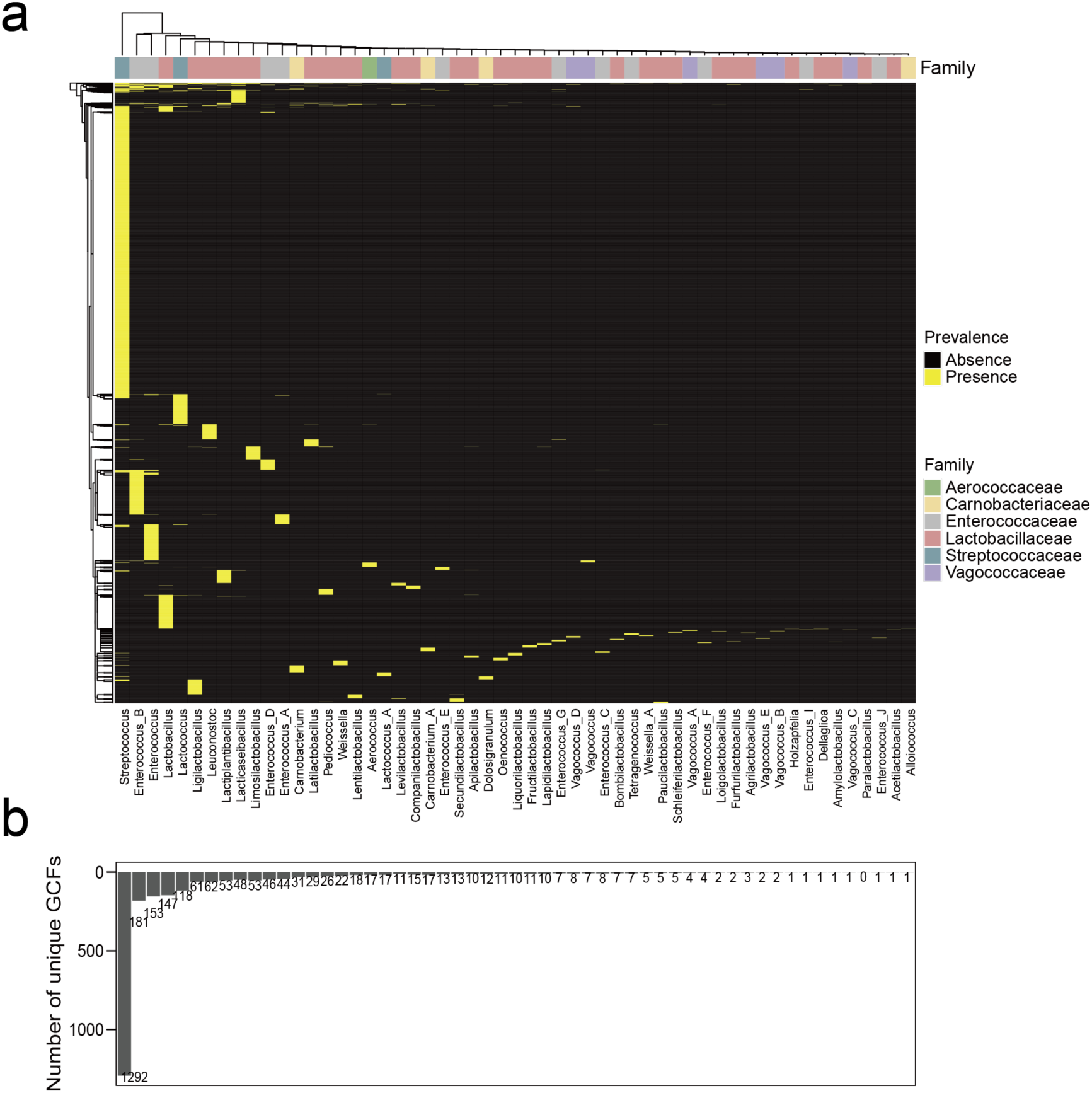
Distribution of 2,849 GCFs. **a**, Distribution of 2,849 GCF in different genera. Each row represents one GCF. The genera of one family do not cluster together, indicating that the GCF between genera varies considerably and does not exhibit phylogeny-relatedness at the genus level. **b**, Number of unique GCFs in each genus.

**Supplementary Figure 9.**
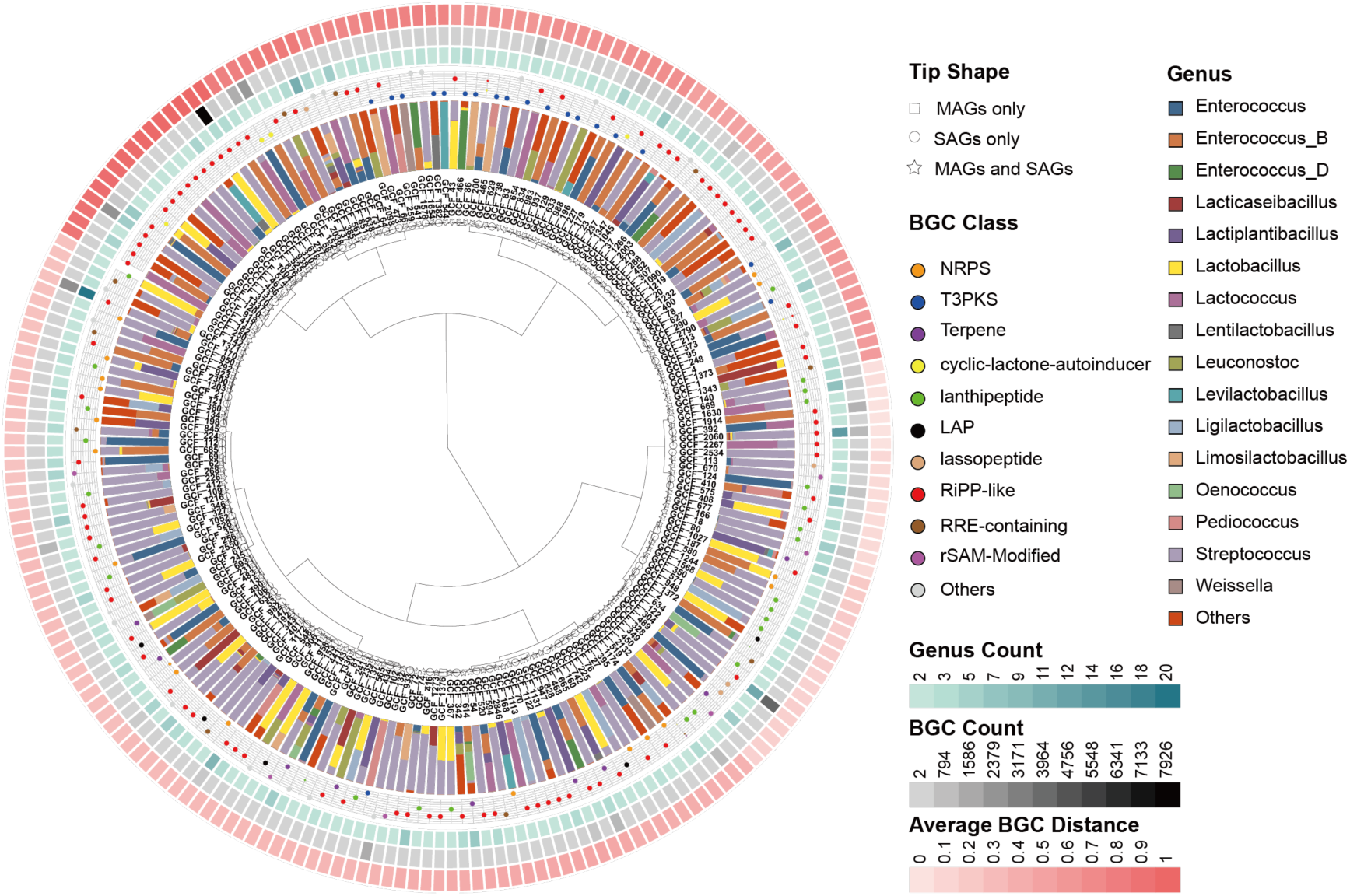
Diversity of 212 cross-genus GCFs. Layers from inner to outer are (1) hierarchical clustering of 212 genera present in >1 genera, based on average BGC distance to known BGCs (experimentally validated BGCs in MIBiG database); (2) the proportion of distinct genera; (3) the percentage of BGC classes in each GCF. Point size is proportionate to the percentage; (4) the number of genera in which the GCF distributes; (5) BGC count in each GCF; (6) average BGC distance to MiBiG BGCs. Among 212 GCFs, the number of GCFs dominated by different classes of BGCs is as follows: NRPS, 17; T3PKS, 19; terpene, 9; cyclic-lactone-autoinducer, 4; lanthipeptide, 4; LAP, 5; lassopeptide, 4; RiPP-like, 88; RRE-containing, 10; rSAM-Modified, 5.

**Supplementary Figure 10.**
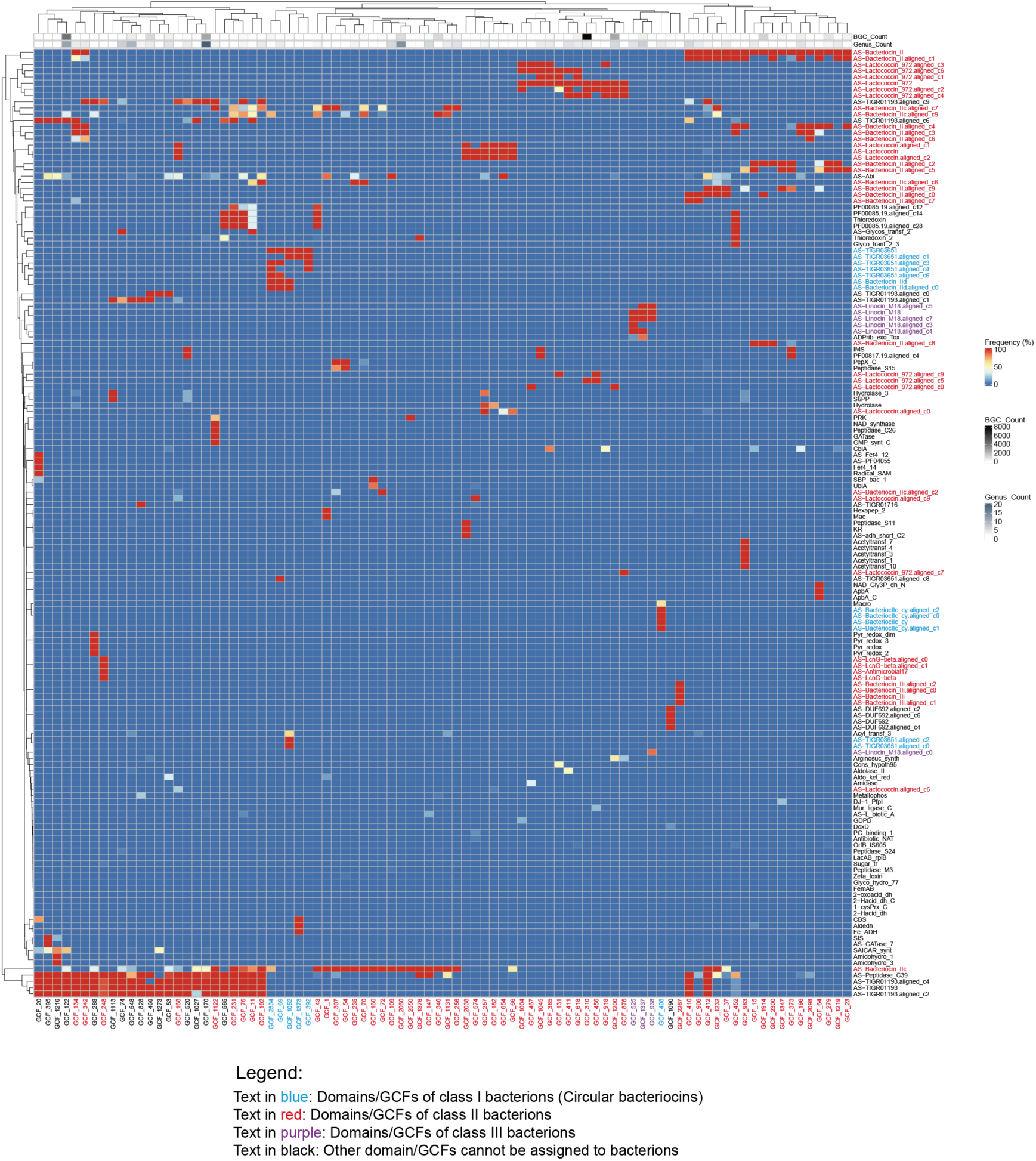
Domain distribution of 88 cross-genus RiPP-like GCFs. The heatmap shows the percentage of predominant BGC features (domains, extracted by BiG-SLiCE) in 88 RiPP-like GCFs, the majority of which are class II bacteriocins (62 GCFs). For each GCF, the top five abundant domains are fetched out, and all abundant domains retrieved are shown on the heatmap. Characteristic domains of class I, class II, and class III bacteriocin are highlighted in blue, red, and purple.

**Supplementary Figure 11.**
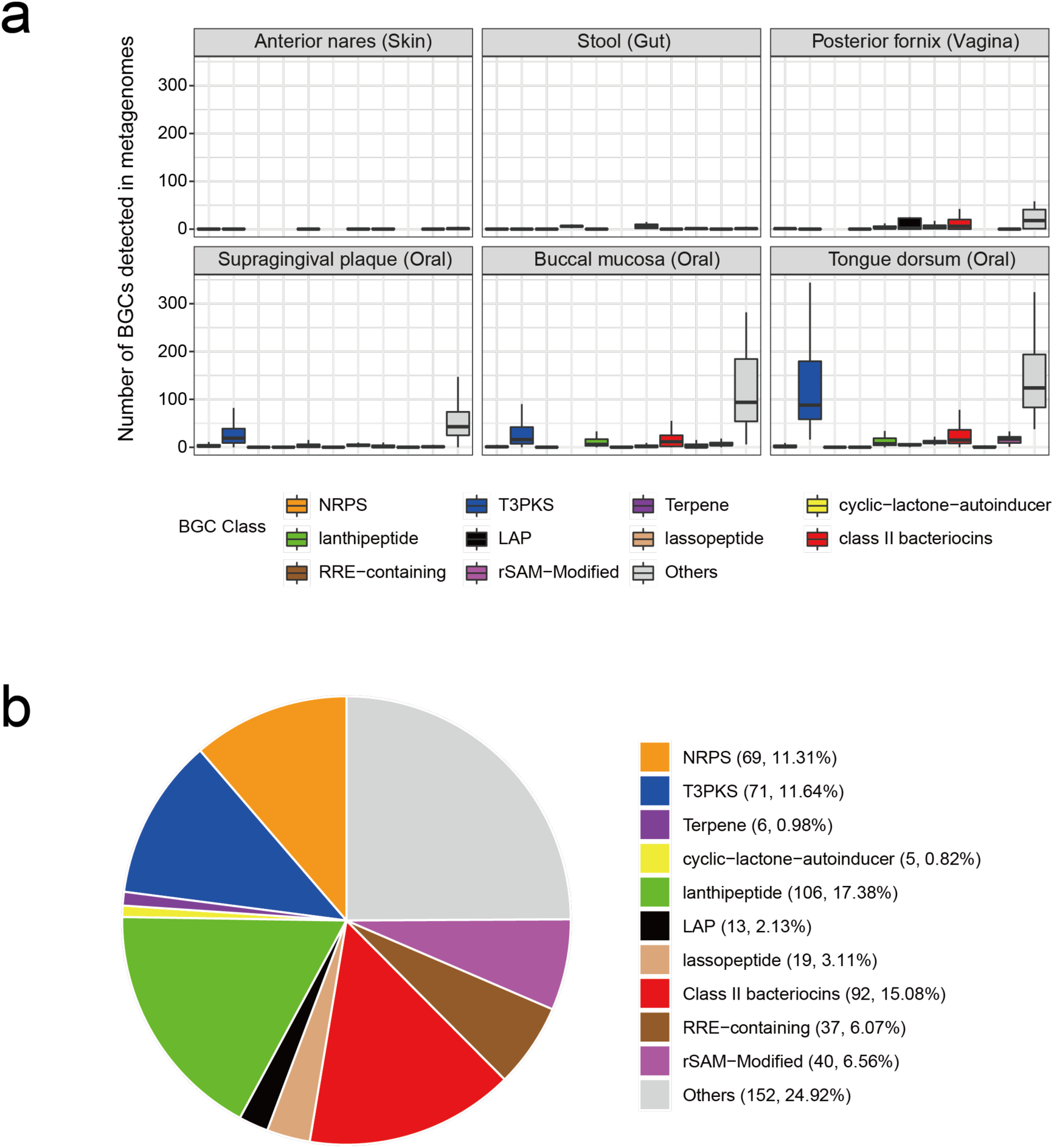
BGCs detected in six body sites. **a,** The box plots show the number of BGCs detected in metagenomes of six body sites. BGCs are stratified as per different classes. BGCs that harbor class II bacteriocins-related domains (shown in Supplementary Figure 9, 10) are grouped into class II bacteriocins. b, The proportion of BGC classes of 610 GCFs detected in six body sites. The numbers in the bracket are the number of GCFs and the corresponding percentage.

**Supplementary Figure 12.**
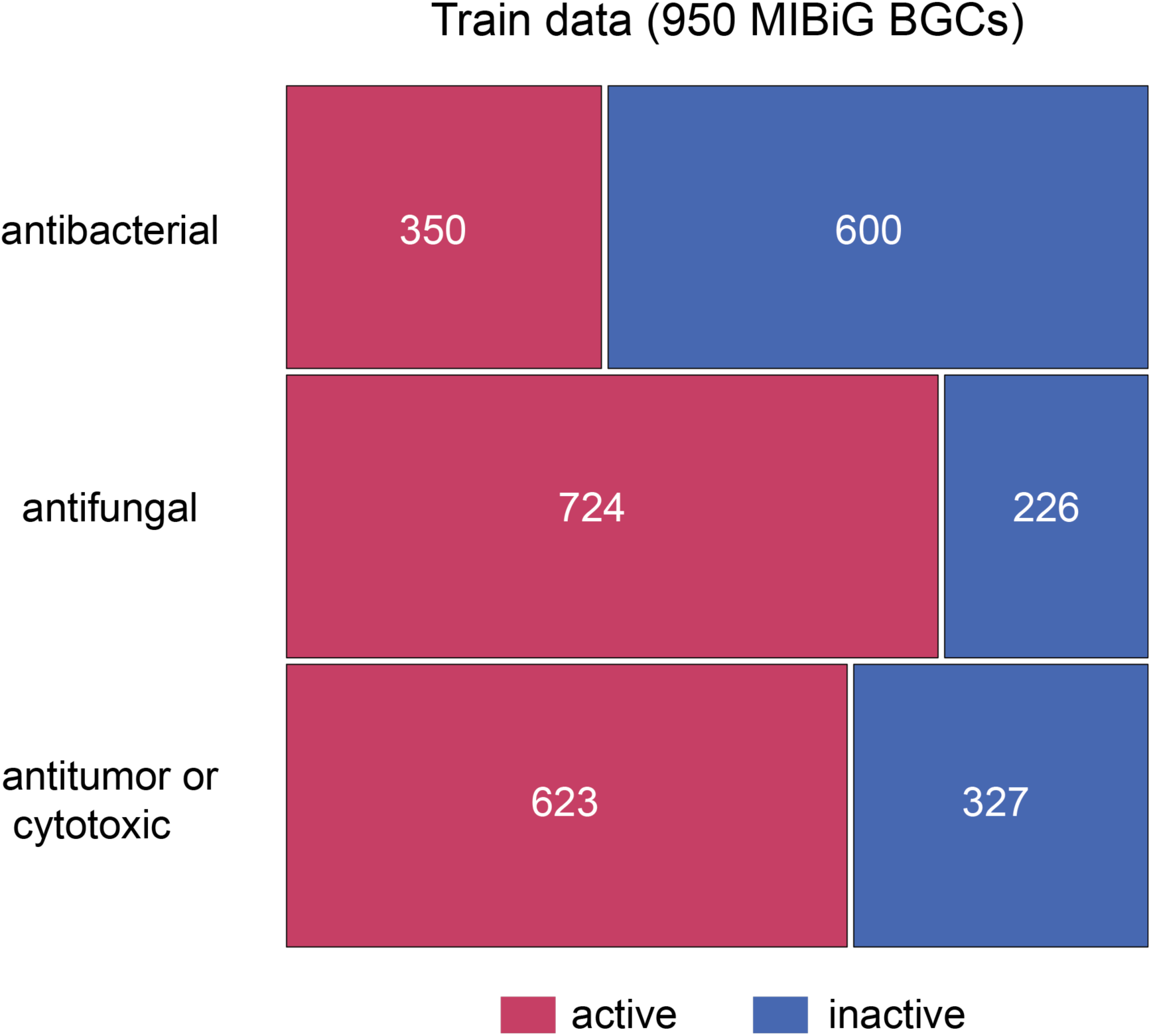
Distribution of reference BGCs with different activities. A total of 950 MIBiG BGCs with known functions were used as the train data set. The number denotes the BGC counts.

**Supplementary Figure 13.**
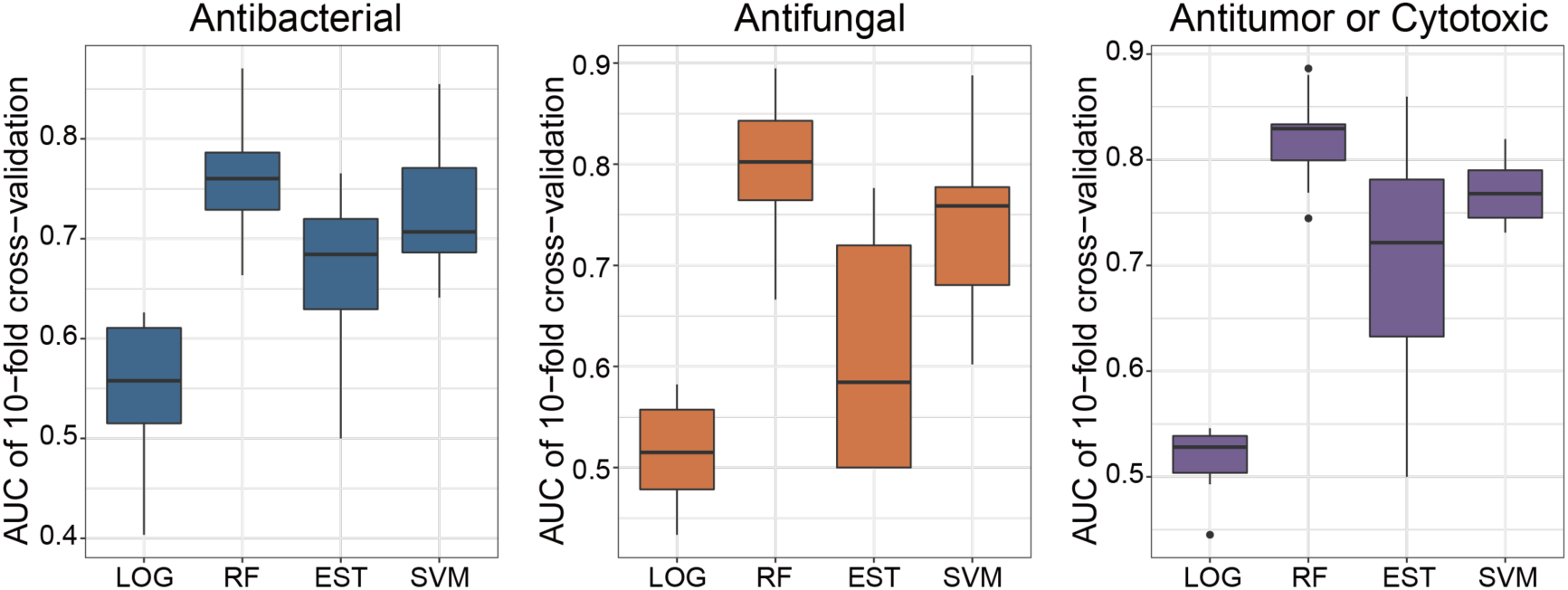
Performances of four classifiers in determining activities of BGC-encoding compounds. The performances were quantified with index AUC [area under the ROC curve (receiver operating characteristic curve)], using 10-fold cross-validation. LOG, logistic regression; RF, random forest; EST, elastic net regression; SVM, support vector machines.

**Supplementary Figure 14.**
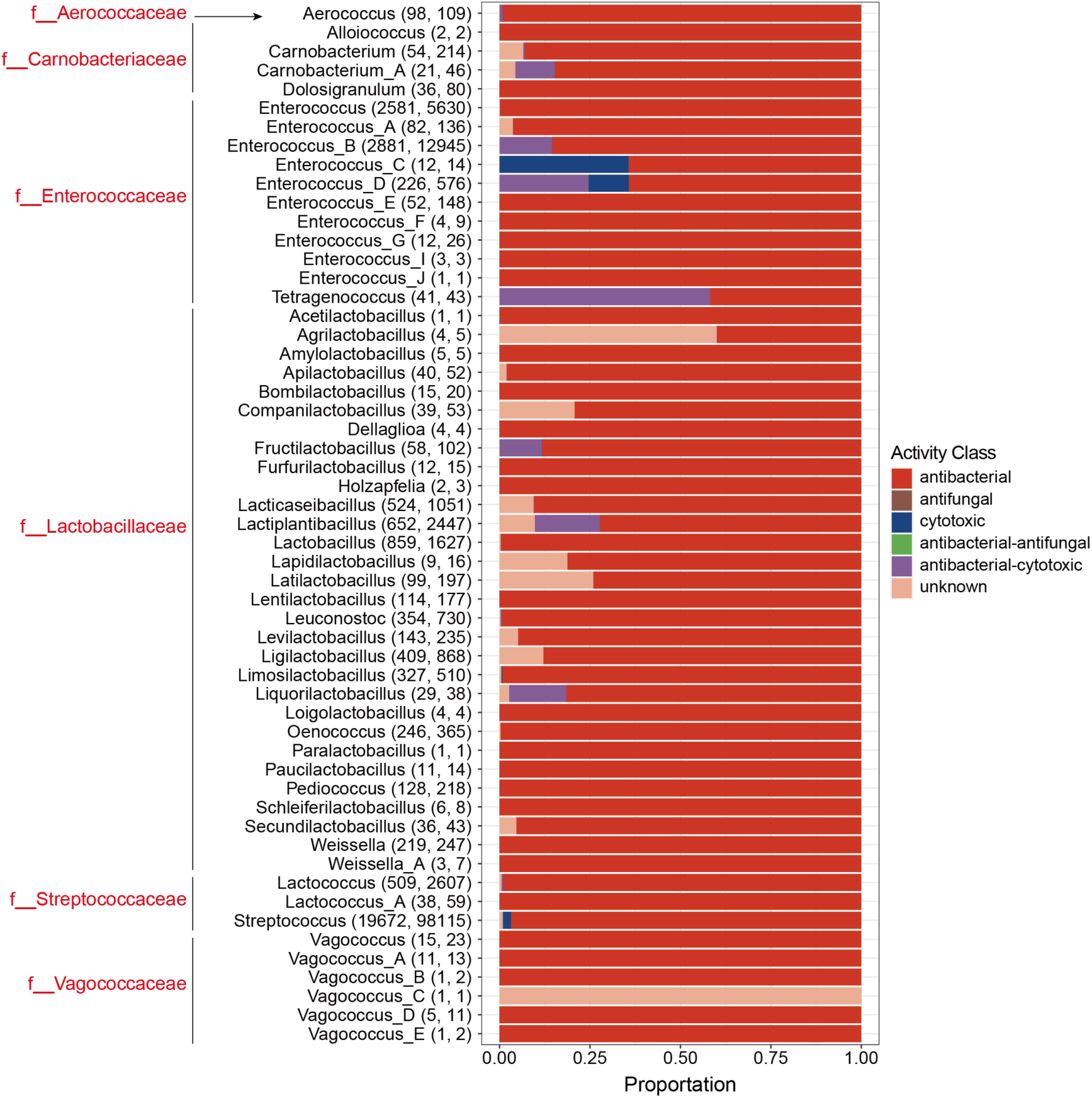
Profile of predicted compound activities of BGCs in LAB genera. The numbers in the bracket are genome count and total BGC count, respectively.

**Supplementary Figure 15.**
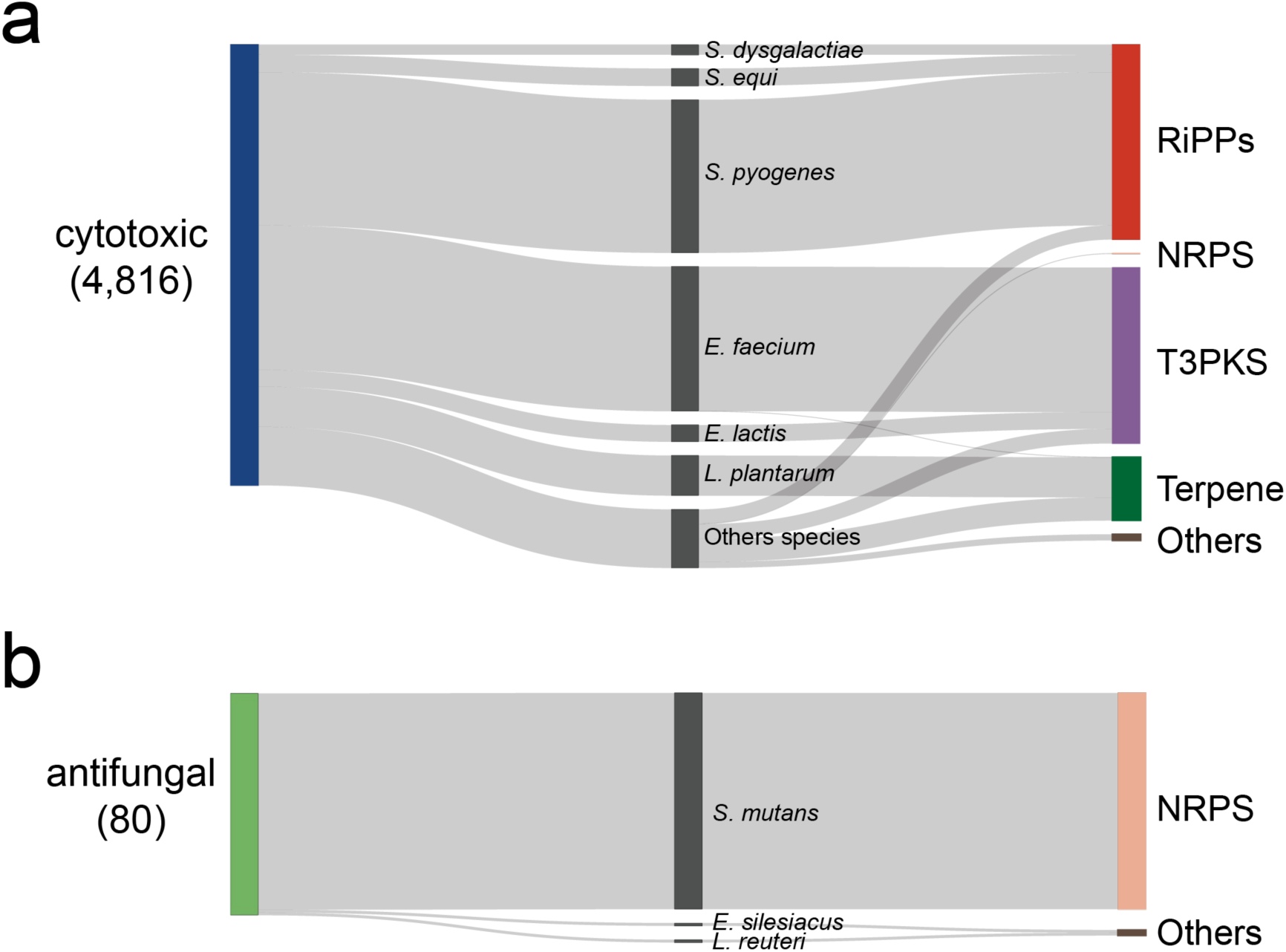
Predicted activity of cytotoxic and antifungal. Sankey diagrams show the BGCs with the predicted activity of cytotoxic (**a**) and antifungal (**b**). The number shown in brackets refers to the BGC count. The full names of bacteria are as follows: *S. dysgalactiae*, *Streptococcus dysgalactiae*; *S. equi*, *Streptococcus equi*; *S. pyogenes*, *Streptococcus pyogenes*; *E. faecium*, *Enterococcus faecium*; *E. lactis*, *Enterococcus lactis*; *L. plantarum*, *Lactiplantibacillus plantarum*; *S. mutans*, *Streptococcus mutans*; *E. silesiacus, Enterococcus silesiacus*; *L. reuteri*, *Limosilactobacillus reuteri*.

**Supplementary Figure 16.**
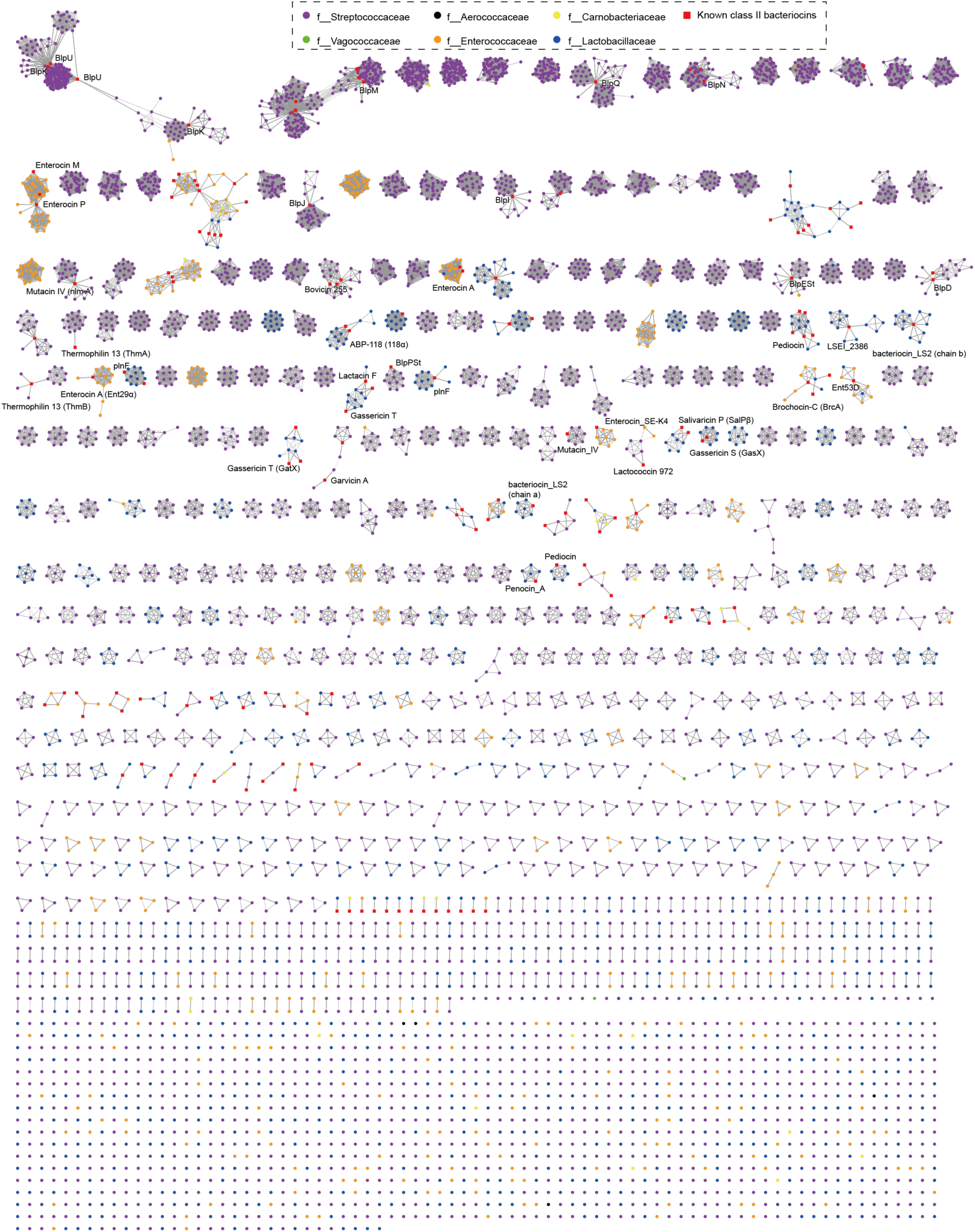
The network of sequence similarity of precursor peptides reveals the huge diversity of putative class II bacteriocins. the 187,649 precursor sequences were de-duplicated to 6,516 sequences, shown in the network. Within each cluster, two sequences with identity > 50% are connected with a line. Precursor sequence and known class II bacteriocins are connected only if the identity > 90% and coverage > 95%. The line width is proportionate to the sequence identity. Each dot represents one precursor sequence, which is further colored according to the LAB family.

**Supplementary Figure 17.**
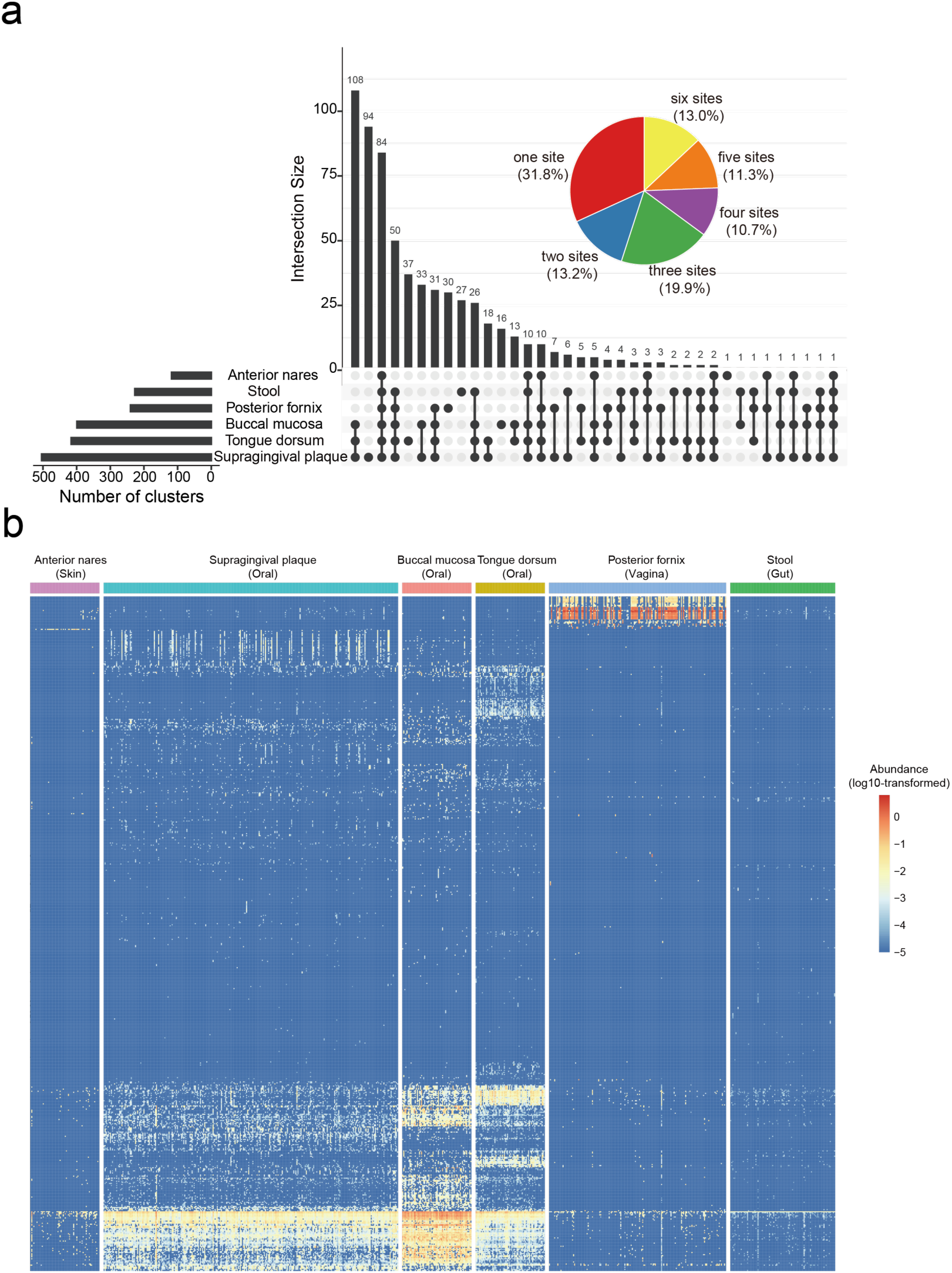
Profile of precursor clusters of class II bacteriocins detected in six body sites. **a**, The intersection of clusters detected in different body sites. The bar plot on the left shows the cluster count in each body site; the bar plot on the top refers to the number of clusters of each intersection. Connecting lines are drawn if an intersection is present in more than one site. The pie chart shows the proportion of 644 clusters detected in different body sites. The corresponding percentages are shown in the brackets. **b**, The heat map shows the profile of 644 clusters in human metagenomes. The heat map reveals different predominant clusters in the vagina and oral cavity.

**Supplementary Figure 18.**
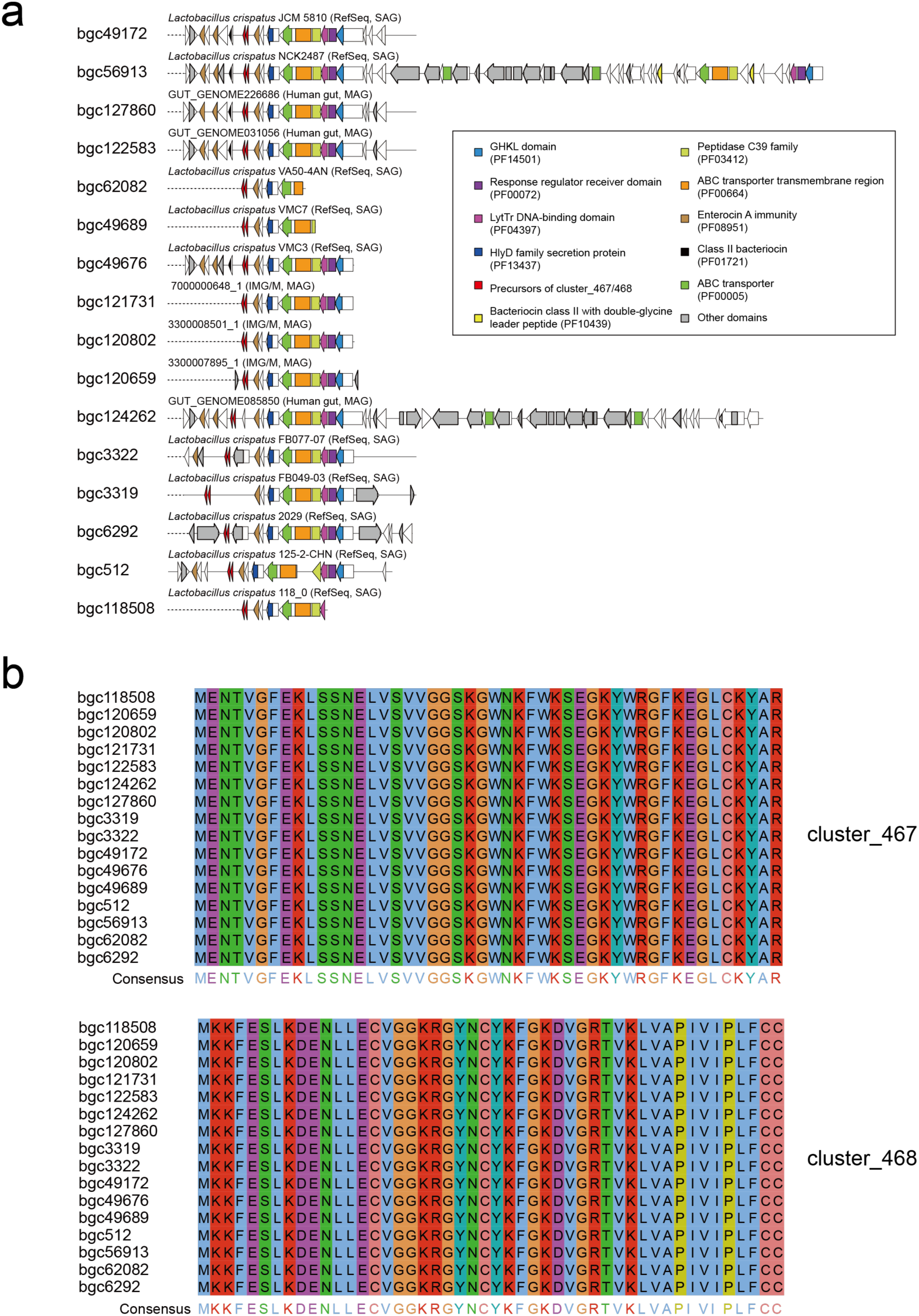
BGCs harboring precursors of cluster_467 and cluster_468. **a**, Gene organizations of 18 BGCs containing putative precursors belonging to cluter_467 and cluster_468. BiG-SCAPE detected the domains. Ten BGCs from isolates deposited in the RefSeq database, three BGCs from human gut metagenomes, and three BGC from metagenomes deposited in IMG/M database were labeled on the top of the BGC architecture. **b**, Precursor peptides corresponding to the 16 BGCs shown in (**a**).

**Supplementary Figure 19.**
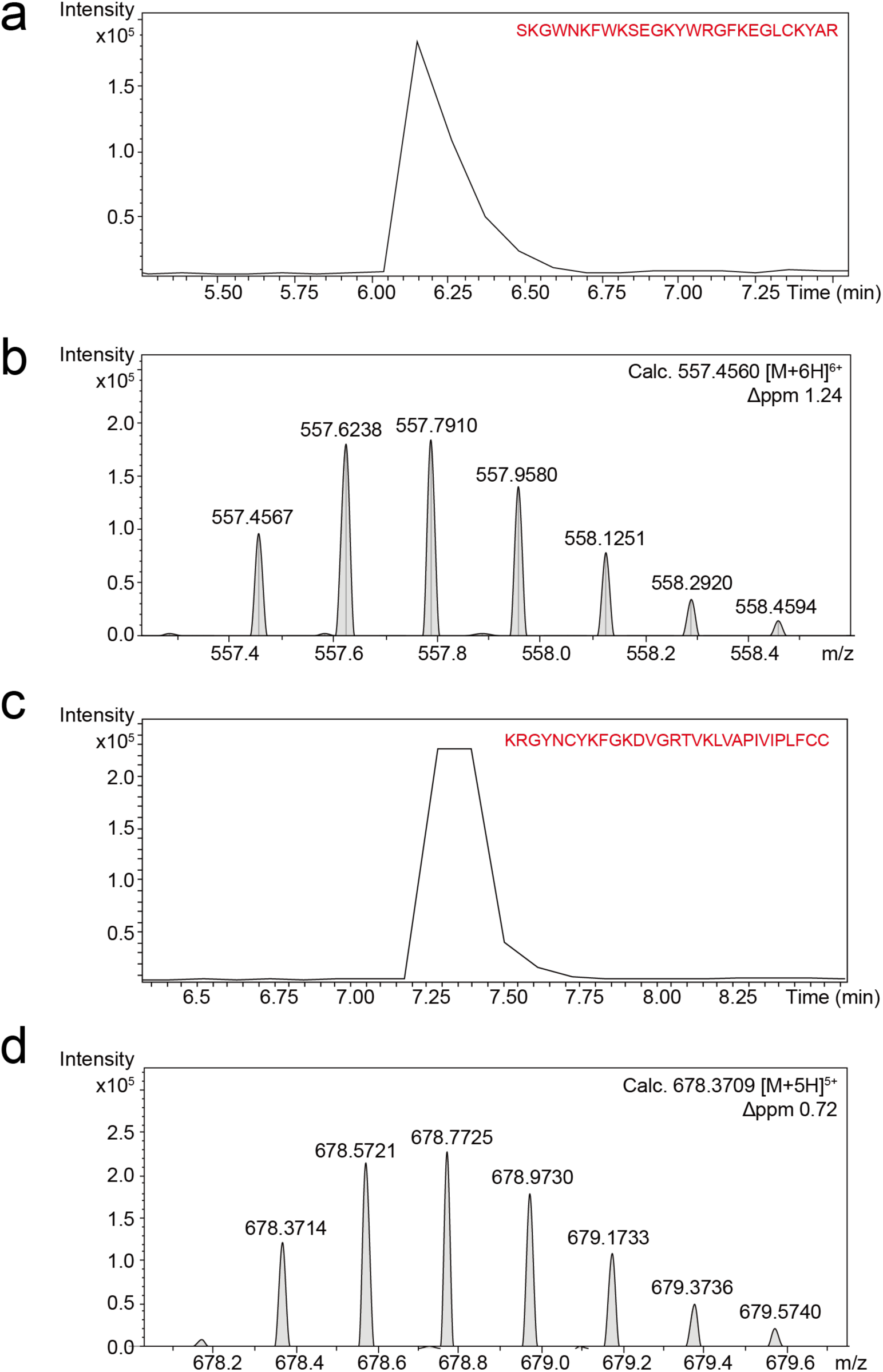
HR-LCMS analysis of synthesized peptides. HR-LCMS analysis of two core peptides is shown here, including the retention time (**a, c**) and their MS1 spectra (**b, d**) for cluster_467 (**a, b**) and cluster_468 (**c, d**).

